# Length estimation of fish detected as non-occluded using a smartphone application and deep learning techniques

**DOI:** 10.1101/2023.03.12.532319

**Authors:** Yasutoki Shibata, Yuka Iwahara, Masahiro Manano, Ayumi Kanaya, Ryota Sone, Satoko Tamura, Naoya Kakuta, Tomoya Nishino, Akira Ishihara, Shungo Kugai

**Affiliations:** Fisheries Resources Institute, National Research and Development Agency, Japan Fisheries Research and Education Agency, Yokohama, Kanagawa, Japan; Marine Resources Research Center, Aichi Fisheries Research Institute, Toyohama, Aichi, Japan; Fisheries Technology Center Sagami Bay Experiment Station Of Kanagawa Prefectural Government, Odawara, Kanagawa, Japan; Computermind Corp., Nishi-Shinjyuku, Tokyo, Japan

**Keywords:** Mask R-CNN, mobile imaging, occlusion, total length composition in catch, stock assessment

## Abstract

Uncertainty in stock assessment can be reduced if accurate and precise length composition of catch is available. Length data are usually manually collected, although this method is costly and time-consuming. Recently, some studies have estimated fish species and length from images using deep learning by installing camera systems in fishing vessels or a fish auction center. Once the deep learning model is properly trained, it does not require expensive and time-consuming manual labor. However, several previous studies have focused on monitoring fishing practices using an electronic monitoring system (EMS); therefore, it is necessary to solve many challenges, such as counting the total number of fish in the catch. In this study, we proposed a new deep learning-based method to estimate fish length using images. Species identification was not performed by the model, and images were taken manually by the measurers; however, length composition was obtained only for non-occluded fish detected by the model. A smartphone application was developed to calculate scale information (cm/pixel) from a known size fish box in fish images, and the Mask R-CNN (Region-based convolutional neural networks) model was trained using 76,161 fish to predict non-occluded fish. Two experiments were conducted to confirm whether the proposed method resulted in errors in the length composition. First, we manually measured the total length (TL) for each of the five fish categories and estimated the TL using deep learning and calculated the bias. Second, multiple fish in a fish box were photographed simultaneously, and the difference between the mean TL estimated from the non-occluded fish and the true TL from all fish was calculated. The results indicated that the biases of all five species categories were within ± 3%. Moreover, the difference was within ± 1.5% regardless of the number of fish in the fish box. In the proposed method, deep learning was used not to replace the measurer but to increase their measurement efficiency. The proposed method is expected to increase opportunities for the application of deep learning-based fish length estimation in areas of research that are different from the scope of conventional EMS.

## 1. Introduction

It is important to accurately estimate current and future population of target fish stocks to maintain maximum sustainable yield (MSY), and population dynamics models, such as age-structured models, have been used to estimate abundance in stock assessments (Ichinokawa et al., 2017; Privitera-Johnson and Punt, 2020). Length-at-age relationships are especially important for these models because the observed fish length compositions are often converted to age compositions using these relationships (Piner et al., 2016). Age composition data are important for estimating recruits, selectivity, fishing and natural mortality rates, and relative stock sizes (Lee et al., 2014; Piner et al., 2016; Shibata et al., 2021). Thus, the length-at-age relationship that has been used to transform length to age is very influential in stock assessments (Wang et al., 2014).

However, obtaining the length composition from landed catch is costly and time-consuming because fish length is usually measured directly by hand (Palmer et al., 2022). In other words, the sample size for fish length of the target species depends on the number of measurers, i.e., the people that measure the fish. This cost often implies reduced sample sizes, which may lead to information loss and negatively affect the accuracy and precision of estimated abundance.

In recent years, some studies have estimated fish length using deep learning methods (Álvarez-Ellacuría et al., 2020; Lekunberri et al., 2022; Ovalle et al., 2022; Palmer et al., 2022). In these studies, cameras were set in fishing vessels or the fish auction centers, and the obtained images of the catch were analyzed using deep learning techniques. Once the deep learning model is properly trained, it does not require expensive and time-consuming manual labor (Lekunberri et al., 2022). The results of these studies could be applied to obtain fish length data and eliminate manual labor dependency. However, three major issues must be considered when installing these camera systems.

First, power supplies and spaces for imaging are needed to setup cameras and secure their field of view. Some large vessels or fish auction centers could carry out length estimation, as in previous studies; however, that is not the case for majority. For example, more than 75% of vessels in America and 80% in Asia, Europe, Africa, and Oceania are less than 12 m long (FAO, 2022 https://www.fao.org/3/cc0461en/online/sofia/2022/fishing-fleet.html). The availability of stable electrical power in small vessels may be limited by the battery capacity when an engine is not running (van Helmond et al., 2020). Although an electronic monitoring system (EMS) for small vessels (i.e., 10–12 m) has been developed using portable power with solar panels, only a single camera can be installed because of space limitations or vulnerability to hidden activity outside the field of view (Bartholomew et al., 2018). Moreover, in some cases, some construction is needed to use a power supply in the fish auction center because power supply is usually not available in these areas (probably to prevent a short circuit due to weathering). In addition, imaging can often be blocked by human movement when belt conveyors carrying fish in a fish auction center or landing fishing port are not installed, which may be the case in most fisheries.

Second, there are difficulties in identifying fish species using deep learning techniques. Identification of Fish species using deep learning techniques involves families and genera (e.g., van Essen et al., 2021). Some examples have been reported as possible situations to identify fish species, although the study is based on a few (ten classes) fish species (Lu et al., 2020), or the surrounding environment of imaging is fixed on the ship (Ovalle et al., 2022).

Finally, occlusion of fish bodies by other fish or objects (i.e., only a part of the fish body is visible in the image) affects identification and length estimation. A previous study estimated the total length through head length using Mask R-CNN (Region-based convolutional neural networks) (He et al., 2017) and weight information (Álvarez-Ellacuría et al., 2020). Another study estimated the total length using a deep learning method corrected using Bayesian estimation (Palmer et al., 2022). A common aspect of both studies was the estimation of the total length as a function of some other information. This method is very useful because it can be used even when fish have been occluded, but some effort may be required because the equation must change depending on various factors, such as fish species and season.

These challenges need to be solved to collect fish length of all fish species using the deep learning method as an alternative to measurers. However, none of them are fundamentally solvable with current equipment and technology. In contrast, it is possible to increase the sample size of fish length data with the burden of measurers decreasing if the purpose of using the deep learning method was changed. Here, image analysis using the deep learning method was used to obtain only the total length composition. The identification of fish species and investigation of discarding or bycatch on commercial vessels using EMS and deep learning methods were not the objective of this study.

We propose a method that combines an application and a deep learning method to assist the measurement of catch length composition data. Measurers identify the fish species, hold a camera in their hands, and directly take photographs of the fish. When photographing, an app called ToroCam (an app to record coordinates of the fish box through the smartphone camera) developed in this study will be used. ToroCam adds the coordinates of the four corners of a box containing fish (fish box) to the JPEG image as pixel values. Subsequently, the value of the cm/pixel can be calculated if the size of the fish box in centimeters is known. After photographing with ToroCam, only non-occluded fish can be detected using deep learning techniques. The length composition in the catch can be obtained by multiplying the value of cm/pixel with the lengths of non-occluded fish in pixels. Although some limitations remain because measurers identify species and take images manually, but the fact that fish length can be collected automatically makes it more efficient because body length information can be typed into a field notebook or computer, and the difficulty of cleaning measuring instruments can be reduced. In addition, continuous security and power supply of the camera are not required. The hardware required is only a smartphone, and a computer for analysis is cheaper than camera systems. Because this is a sampling rather than a full count measurement, fish length extracted from only non-occluded fish can be used for stock assessment when the occlusion is random for actual fish size.

The purpose of this study is to develop a method where measurers carry out direct imaging (mobile imaging) with a portable tool, such as a smartphone, even under unsteady imaging conditions, and obtain the total length of non-occluded fish using deep learning techniques. This method will enable the determination of fish length composition by image analysis, even in situations where the methods of fish image analysis developed based on conventional EMS have not been targeted.

## 2. Materials and Methods

### 2.1 Image acquisition for learning

We obtained 8,087 fish images from a fish market, two fishing ports, a bottom trawl vessel, and a research vessel (R/V): Matsuura fish market, Odawara and Toyohama fishing ports, bottom trawl vessel (Horyo-maru No. 18), and R/V Kaiyo-maru No. 5 (Table 1). Horyo-maru No. 18 and R/V Kaiyo-maru No. 5 operated commercial bottom trawling and a scientific bottom trawling survey, respectively, in the southern Hokkaido area (Fig. 1). Fish images were either captured with a fixed camera or manually captured by researchers. The images were collected using a camera mounted directly above a conveyor belt on the Matsuura fish market, Odawara fishing port, and the bottom trawl vessel or the sorting platform of R/V Kaiyo-maru No. 5 (Fig. 2a-d). In addition, at the Matsuura fish market, Odawara, and Toyohama fishing ports, researchers used a camera and directly photographed landed fish (Fig. 2e-g). The images were captured at 3:00–17:00 at the Matsuura fish market, 4:00–7:00 at the Odawara fishing port, 16:00–17:00 at the Toyohama fishing port, 7:00–19:00 at the Horyo-maru No. 5, and 7:30–17:00 at the R/V Kaiyo-maru No. 5.

**Fig. 1.**
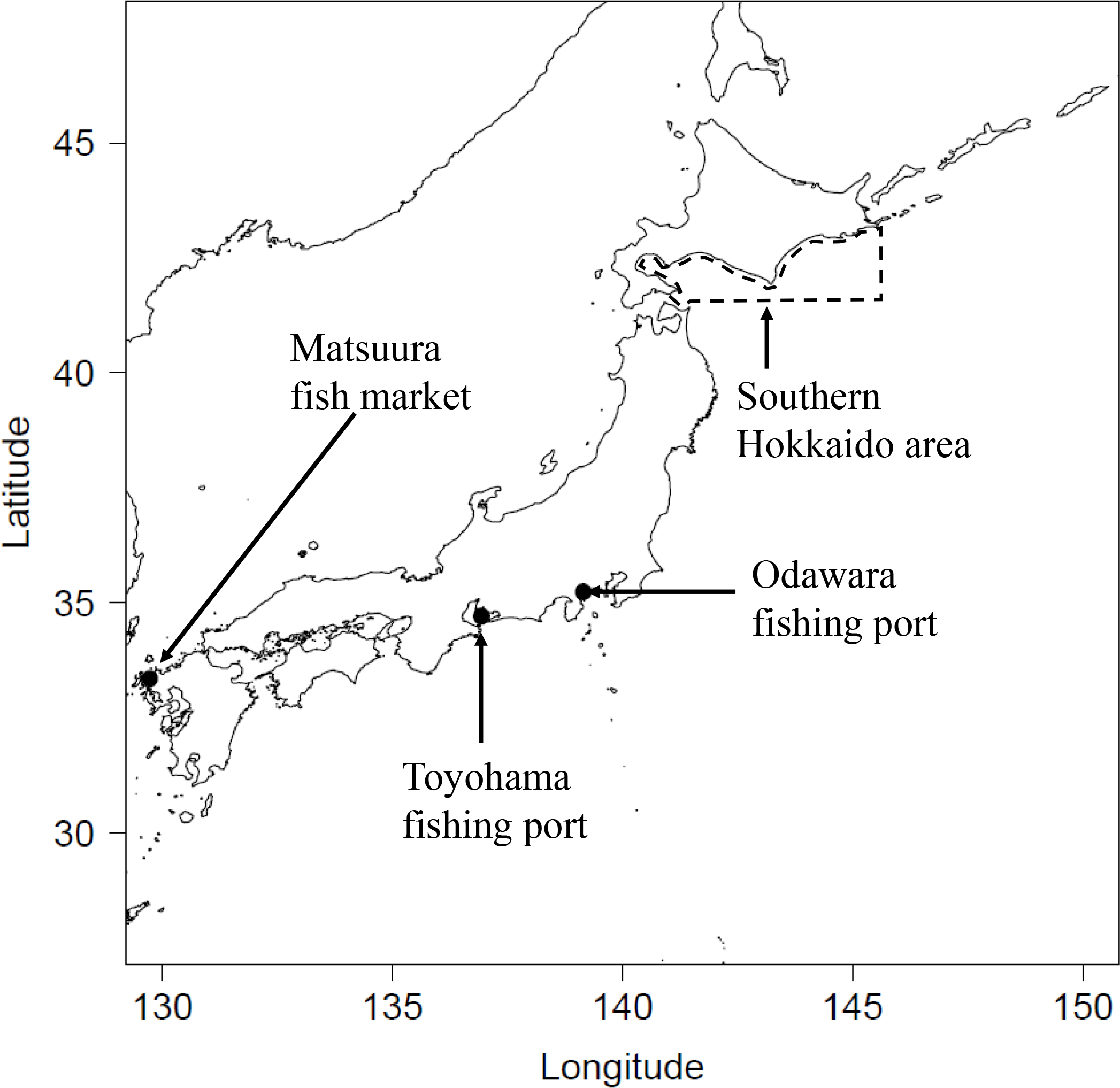
Positions of the Matsuura fishing market, Odawara and Toyohama fishing ports, and southern Hokkaido area where Kaiyo-maru No.5 and Horyo-maru No. 18 were operated.

**Fig. 2.**
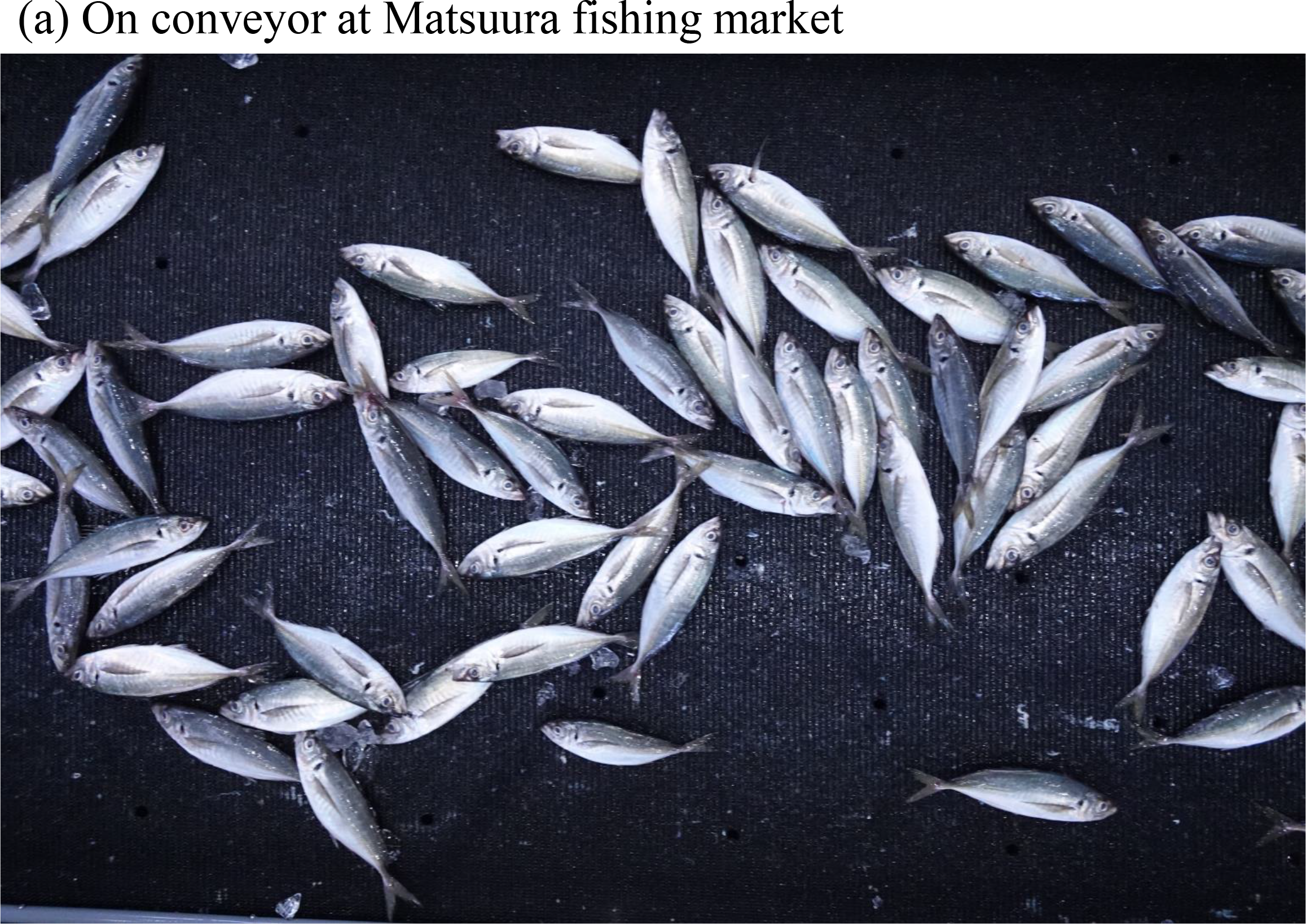
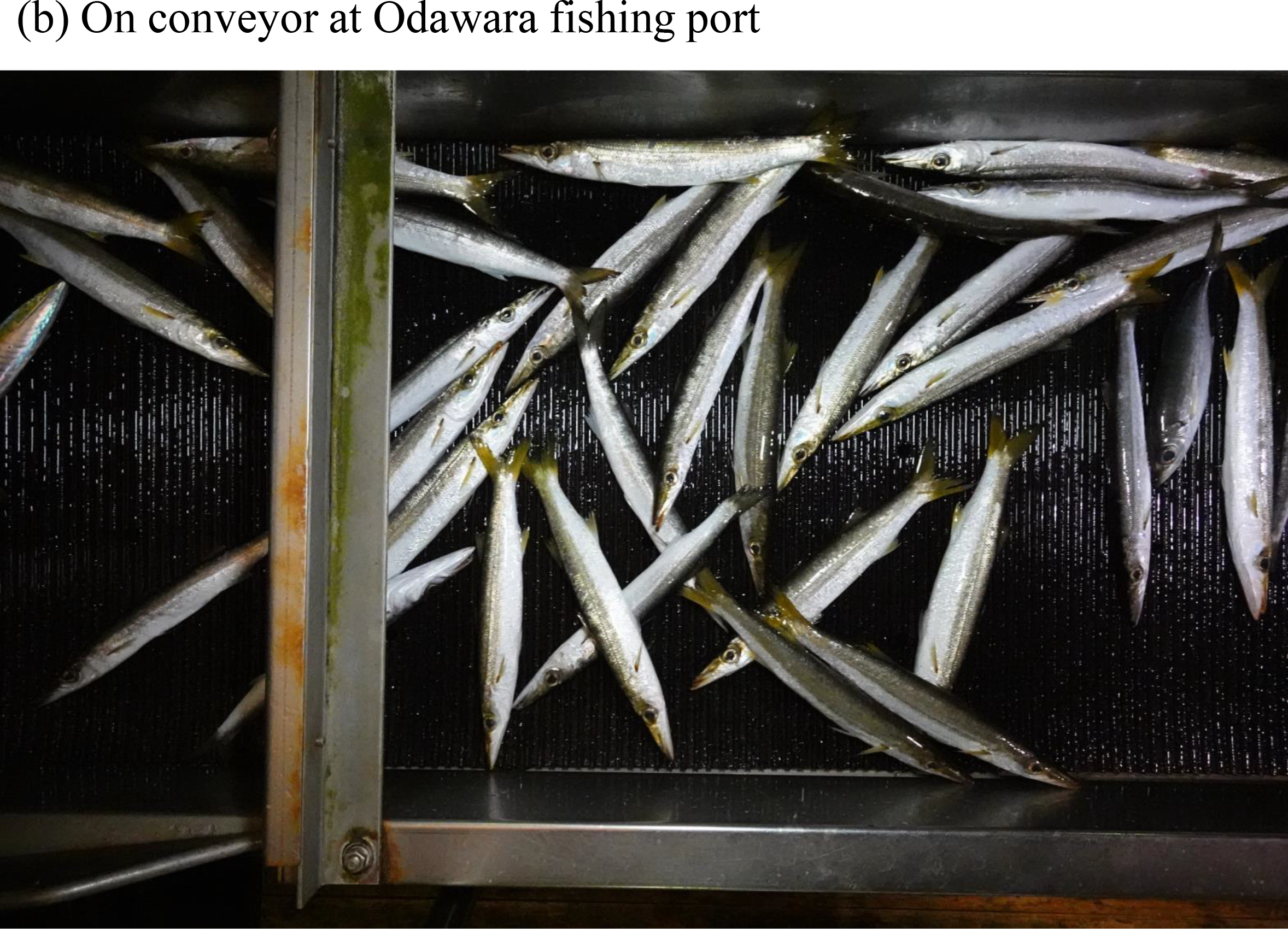
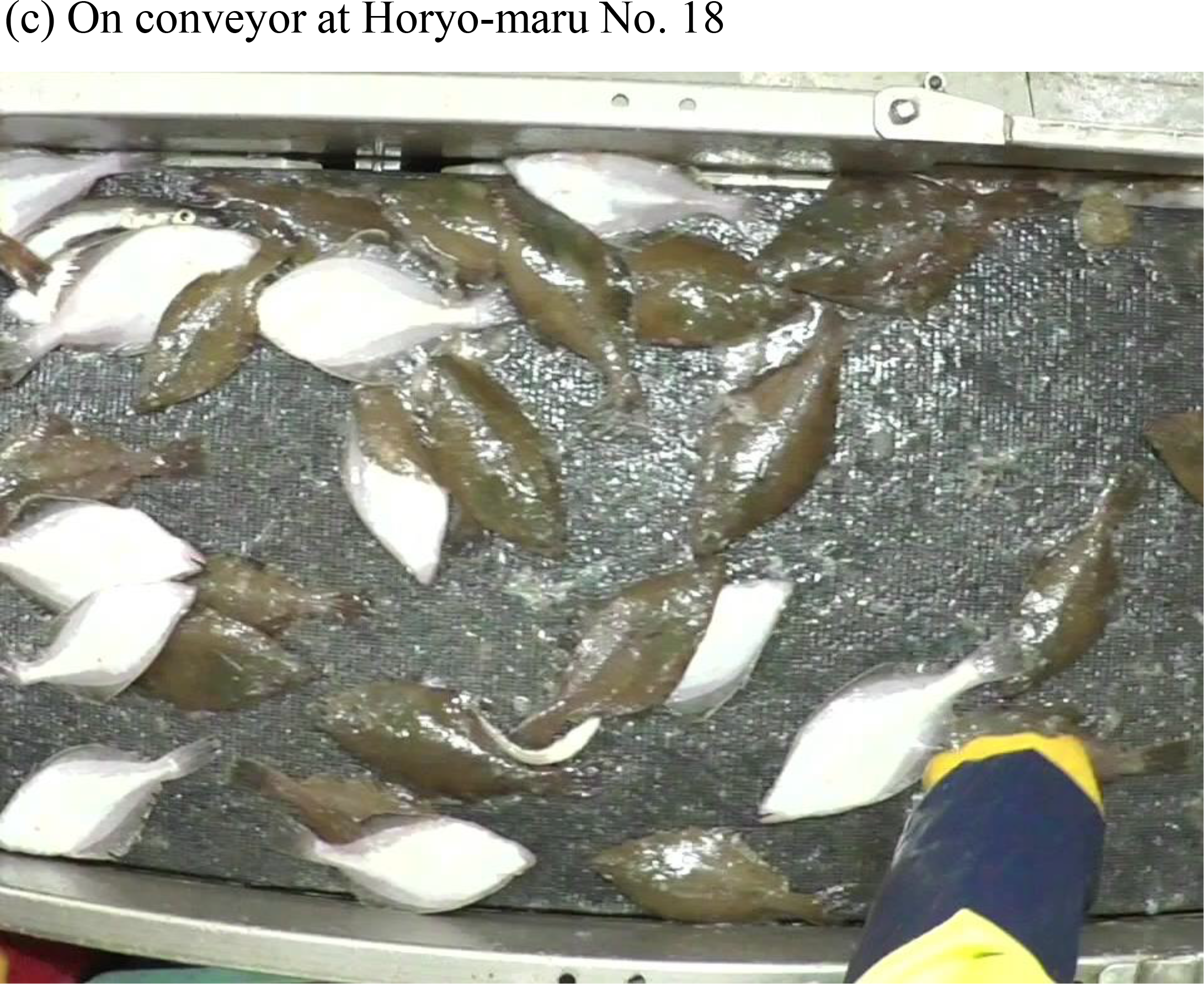
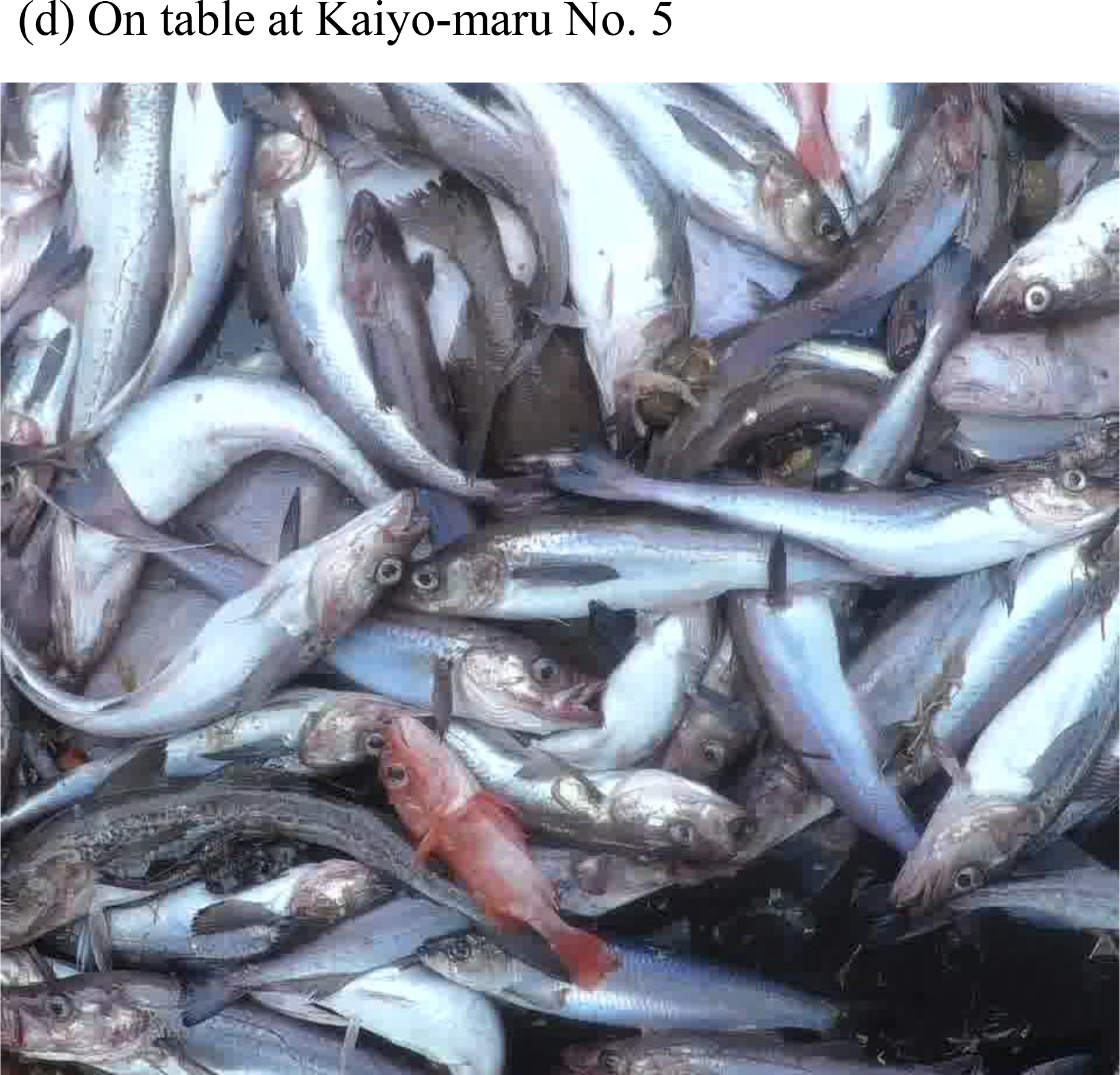
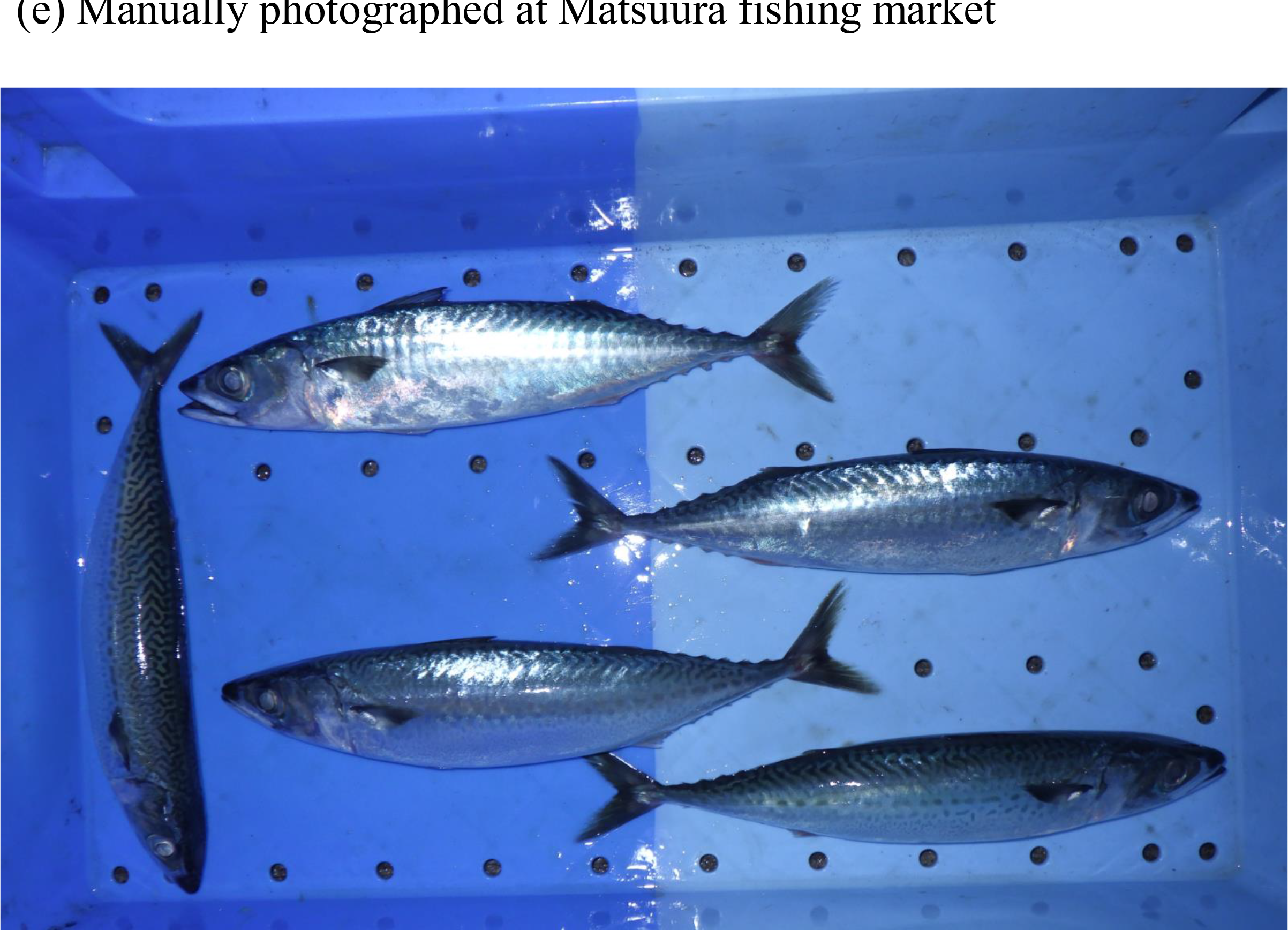
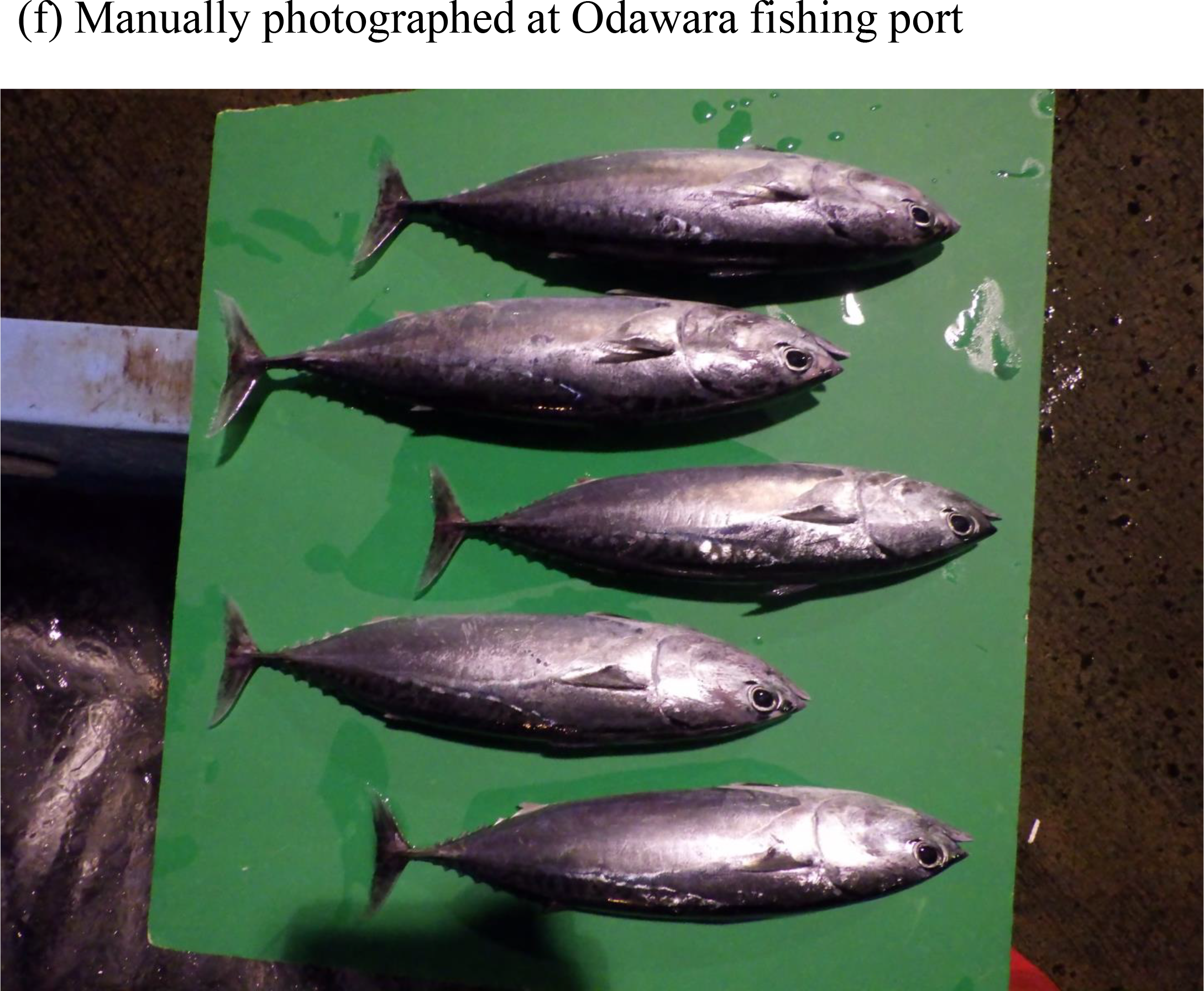
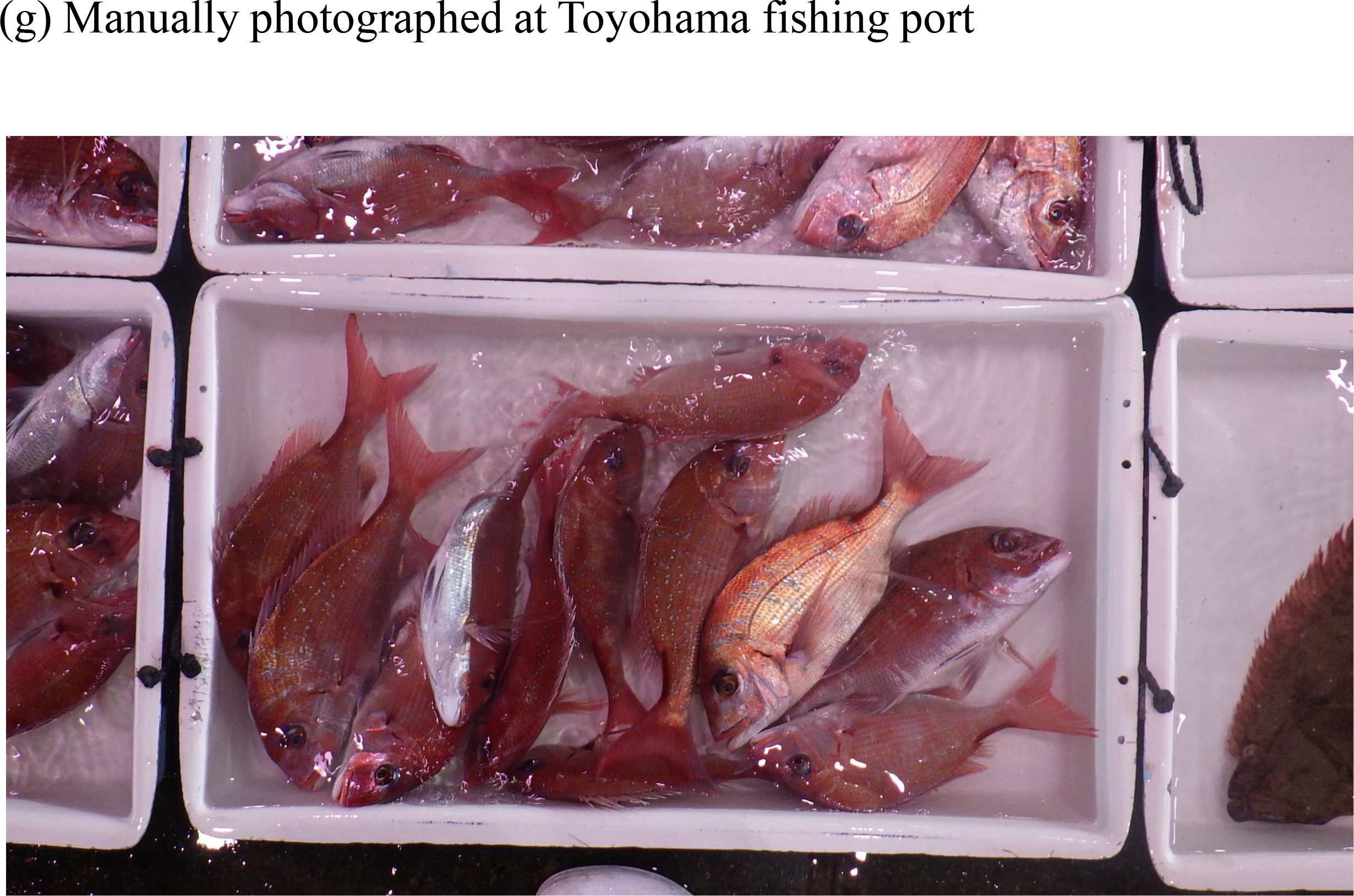
Examples of images that are used for this analysis; (a) images from a camera set up near a conveyor belt at Matsuura fishing market, (b) Odawara fishing port, (c) Horyo-maru No. 18, (d) images from a camera set up near a sorting table on Kaiyo-maru No. 5, (e) a manually photographed image at Matsuura fishing market, (f) Odawara fishing port and (g) Toyohama fishing port.

**Table 1.** The number of images by image size and obtained places that were used in this analysis.

|  | Matuura<br>fish market | Odawara<br>fishing port | Toyohama<br>fishing port | Kaiyo-maru<br>No. 5 | Horyo-maru<br>No.18 | Total |
| --- | --- | --- | --- | --- | --- | --- |
| 4800x3200 | 119 | 3,891 |  |  |  | 4,010 |
| 2048x1536 |  | 1,769 |  |  |  | 1,769 |
| 2704x1520 |  | 1,229 |  |  |  | 1,229 |
| 1920x1080 |  |  | 360 | 55 |  | 415 |
| 5184x3888 | 193 | 115 | 80 |  |  | 388 |
| 960x1080 |  |  |  | 41 | 61 | 102 |
| 3072x1728 |  |  | 101 |  |  | 101 |
| 4608x3456 |  |  | 40 |  |  | 40 |
| 424x480 |  |  |  |  | 27 | 27 |
| 640x720 |  |  |  |  | 6 | 6 |
| Total | 312 | 7,004 | 581 | 96 | 94 | 8,087 |

The image sizes were as follows: 4800 × 3200, 2048 × 1536, 2704 × 1520, 1920 × 1080, 5184 × 3888, 960 × 1080, 3072 × 1728, 4608 × 3456, 424 × 480, and 640 × 720. The images were taken from August 2020 to December 2021, whereas those from Horyo-maru No. 18 were obtained from December 2015 to February 2016. Because the images obtained from Horyo-maru No. 18 and Kaiyo-maru No. 5 were taken in fish-eye mode, 25% of the left and right side of the images were removed, and the remaining 50% in the middle was extracted as the region of interest and used for the following analysis.

### 2.2 Annotation and classes

All fish in the images were annotated using instance segmentation (n = 76,161). Because fish identification was not the target of this study, detailed names of fish species and their number of annotations are shown in Supplementary Table 1. All fish bodies, including fins and spines, were annotated. The mantle was annotated for squids.

Exposure of the fish body was annotated as “*exposure”*. The label was classified on two scales as “F-100” and “F-other.” For example, if the fish body is not occluded by another fish body or object, the label *exposure* would be “F-100.” If the body’s visibility is reduced from 99 to 1% of its whole body because of occlusion, then *exposure* was “F-other.” The numbers for each category are shown in Table 2. In addition, if fish could not be annotated one by one and fish species could not be determined or the fish was outside the conveyor, they were labeled as “Non-target” and those were annotated in a same polygon (Fig. 3b). A difference between “Non-target” and “F-other” is that the former does not provide instance segmentation for each individual and does not contain information that can be used for species identification, while the latter can be used even if it is occluded. This difference may not be very useful in this study because we did not perform species identification. However, we anticipate that it will be useful to keep the two separate considering future studies. If a fish is detected and identified as “F-100” by a deep learning model in inference, the fish can be used to obtain the total length composition.

**Fig. 3.**
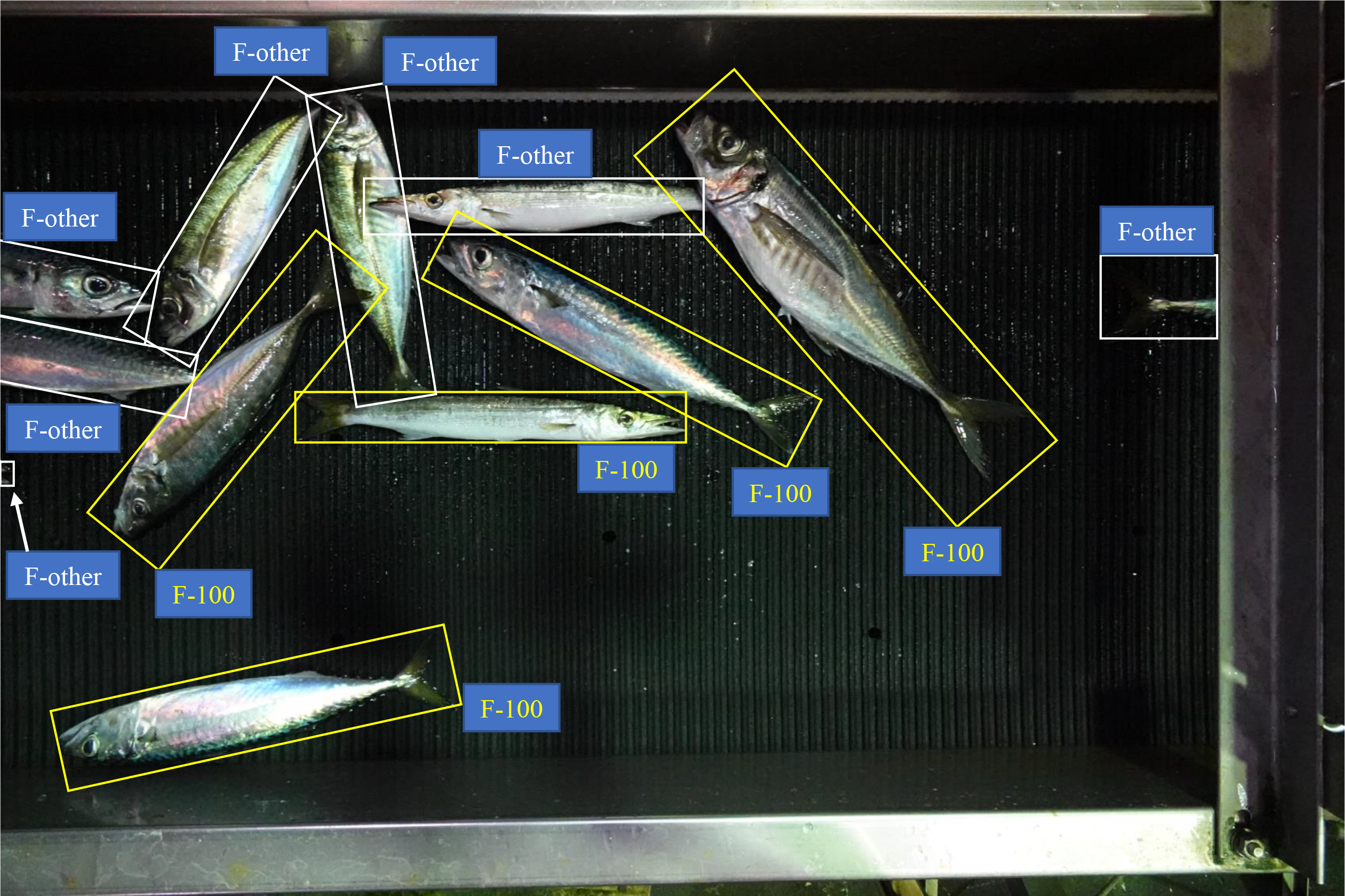
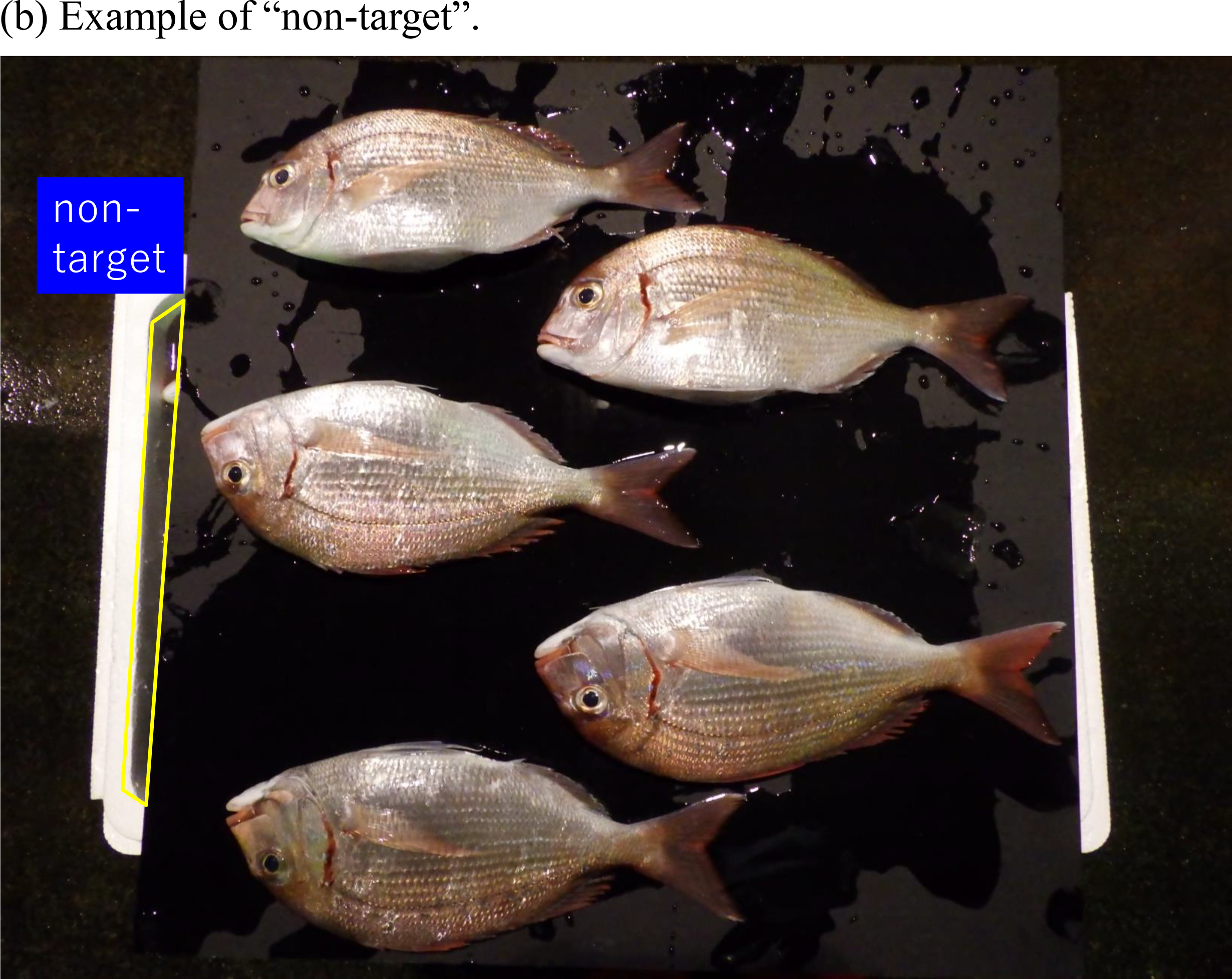
(a) Examples of *exposure* where mask area is not drawn to keep visibility of original fish body. An individual fish annotated as “F-100” is shown in the yellow colored bounded boxes and text, whereas those of “F-other” are shown in white. (b) Examples of the label “Non-target” where the polygon area includes parts of fish body but cannot identify their species and number.

**Table 2.** The number of images and individuals or objects that are classified as “F-100”, “F-other”, and “Non-target” by obtained places. These are used for training data.

|  | Matuura<br>fish market | Odawara<br>fishing port | Toyohama<br>fishing port | Kaiyo-maru<br>No. 5 | Horyo-maru<br>No.18 | Total |
| --- | --- | --- | --- | --- | --- | --- |
| Images | 312 | 7,004 | 581 | 96 | 94 | 8,087 |
| F-100 | 2,407 | 17,468 | 571 | 240 | 159 | 20,845 |
| F-other | 4,675 | 39,753 | 6,504 | 1,209 | 3,175 | 55,316 |
| Non-target | 0 | 1,535 | 149 | 28 | 76 | 1,788 |

### 2.3 Training and validation

Eighty percent of the annotated images were used for training, and the remaining 10% were used for validation. The remaining 10% was used as the test data to calculate the confusion matrix. During training, the image size was resized to 800 × 600 pixels, flipped upside down with 50% probability, and data expansion was performed. A PyTorch library (Paszke et al., 2019) included in Python was used, and the estimation was performed using Mask R-CNN (He et al., 2017). The development environment was PyTorch 1.7.1, Python3.7, and a GPU NVIDIA 3800. The number of epochs was fixed at 65, batch size was 4, and IoU (Intersection over Union) was 0.5. ResNet-50-FPN was used as the backbone; the model classified only three classes: “F-100,” “F-other” and “Non-target.” Here, only those with a probability of 0.8 or higher were used to exclude ambiguous objects. The value of the loss function was calculated from the data used for validation, and the weight parameters were adopted when the value was the smallest.

### 2.4 Experiments and inference

Two experiments were conducted from September 2022 to January 2023 to test the performance of estimating fish length using “F-100” with ToroCam. A smartphone (Sony, Xperia 5III, Android12, 0.82 × 6.8 × 15.7 cm) was used to carry out the two experiments and the size of image was fixed at 3024 × 2268. These image sets were not included in the training or validation datasets.

#### 2.4.1 Experiment I. Difference between true and estimated length

The purpose of this experiment was to evaluate the difference in the total length between the measured and estimated individuals. Each fish was photographed and measured individually. At the Matsuura fish Market and Odawara fishing port, the landed catch was randomly sampled and photographed after measuring the total length of each fish. The measured total length was considered the true value. The shooting time was within the same range as that when the training data were obtained. The species of fish photographed were mackerels (*Scomber japonicus* and *Scomber australasicus*), Japanese jack mackerels (*Trachurus japonicus*), Japanese sardines (*Sardinops melanostictus*), red barracudas (*Sphyraena pinguis*), and bullet tuna (*Auxis rochei*). Bullet tuna were only sampled at the Odawara fishing port, and the other five species were sampled from the Matsuura fish market. *Scomber japonicus* and *Scomber australasicus* were difficult to identify in the field survey; therefore, these two species were treated under the same name category. Hereafter, when referring to both simultaneously, they are denoted as mackerels. The sample size (*N*) of mackerels was 150, that of bullet tunas was 100, and that of the remaining species was 50.

The fish were photographed in a blue colored fish box (73 × 40.5 cm), although this background color beneath the fish was not included in the training data. The fish were photographed using ToroCam, a smartphone application (only for Android OS and the app is now ready to be registered on Google Play) that displays a rectangle on the smartphone screen and assigns the coordinates of the rectangle to the photographed JPEG image (Fig. 4). In other words, the photographer must visually align the rectangle on the screen with the corners of the fish box. Not only the coordinates, but also the name of the fish, the true length of the fish box, and the location of the photograph can be recorded automatically once declared at the time of shooting in the comment section of the JPEG image.

**Fig. 4.**
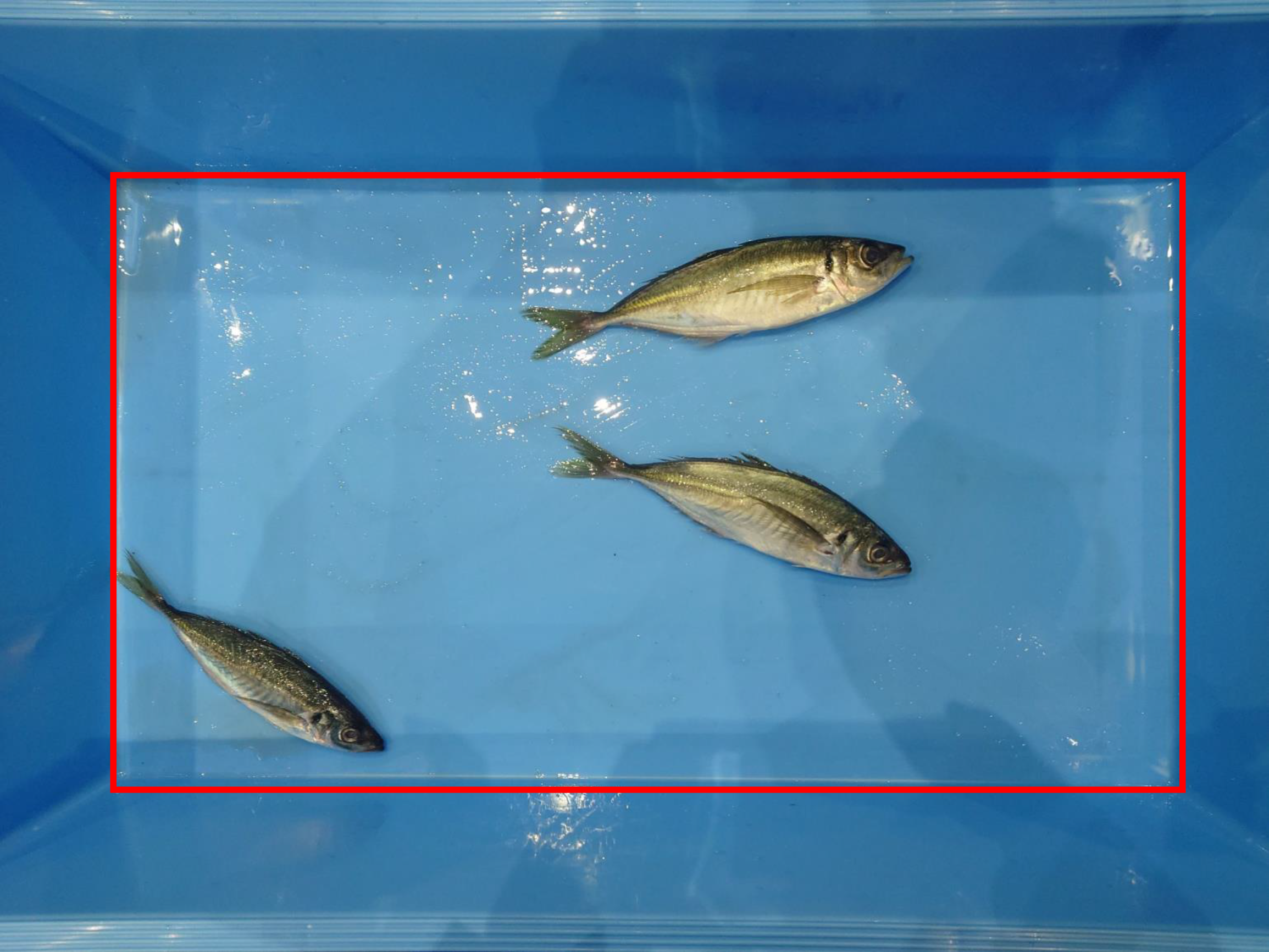
Example of an actual screen through ToroCam. The red rectangle is shown in the camera screen to manually align the corners of rectangle with the corners of the fish box.

ToroCam displays a rectangle on the camera screen of the smartphone, and after aligning the rectangle with the four corners of the fish box containing the fish, the photographer takes a picture. Because the distances between the four corners of the fish box are easily known, the scale information (cm/pixel) was obtained by calculating the number of pixels in the image between the coordinates of the four corners. The fish determined as “F-100” in the image at the time of inference was enclosed by a rectangle, and the number of pixels at long side was multiplied by the calculated cm/pixel value and was converted to the total length. The relative differences *d̂*_*n*_ between the *n*th observed total length *l_n_* and the calculated total length *l̂*_*n*_ and relative bias (*B̂*) were calculated for each fish species as follows:

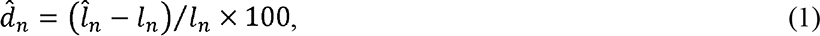

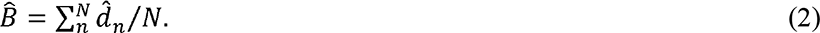

#### 2.4.2 Experiment II: Difference with increasing individuals in a fish box

The purpose of this experiment was to evaluate the degree of change in the difference between the estimated and true length composition when fish in a fish box were increased. The experiment was conducted at the Matsuura fishing market using two fish categories: Japanese jack mackerels and mackerels. Fifty fish were prepared for each category. Fish were added individually to the blue fish box and photographed until *N* = 10 (*N*=1, 2, …, 10). Thereafter, fish were added twice and photographed (*N*=12, 14,…,50). Here, we photographed three shots (*i*=1,2,3) of each *N* with the fish randomly stirred by hand each time. In other words, the same fish were photographed three times, although their positions and degrees of occlusion differed each time. The experiment was conducted over two days, on January 17 and 18. Thus, 180 ((10 + 20) × 3 × 2) photographs were taken for each fish category.

The *n*th fish (*n*=1,…,*N*) in the box were identified as three classes or missed to be detected by the deep learning model: “F-100” (*c*=1), “F-other” (*c*=2), “Non-target” or “not-detected,” although “Non-target” fish were not included in this experiment and “not-detected” fish could not be counted. If the occluded area is sufficiently large, the underlying fish are not visible, and the fish may not be detected. The number of detected fish (*N̂*_*N,c|i*_) as either “F-100” (*N̂*_*N,c = |i*_) or “F-other” (*N̂*_*N,c=2|i*_) was counted, where the value of *N̂*_*N,c|i*_ was changed for each shot *i* even if *N* was the same because the degree of occlusion was changed by hand. The detected rates of only non-occluded individuals (*R̂*_*1,N,i*_) and both non-occluded and occluded individuals (*R̂*_*2,N,i*_) were calculated for every shot *i* under the true number *N* using the following equation:

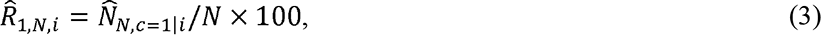

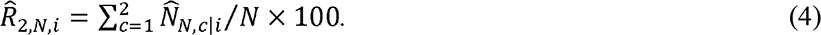

The total length of *n*th individual identified as either *c* = 1 or *c* = 2 for every shot *i* under the true number in the fish box *N*(*l̂*_*n,N,c|i*_) was estimated using ToroCam and deep learning methods. The method used to estimate the total length was the same as that in Experiment 1, although the estimates were obtained for *c*=1 and *c*=2. The mean total length of both non-occluded individuals (*L̂*_*1,N,i*_) and all detected individuals *L̂*_*2,N,i*_ was calculated for every shot *i* and *N* as follows:

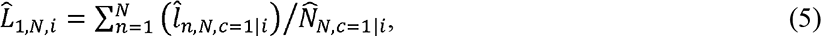

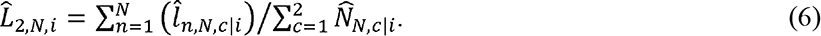

Note that the *n*th estimated total length is not added if the individual is identified as *c*=2 in Eq. (5) but Eq. (6). Finally, a relative difference at each shot *i* of only non-occluded individuals (*D̂*_1,*N, i*_) and all detected individuals (*D̂*_2,*N, i*_) under the true fish number *N* was calculated using the following equation:

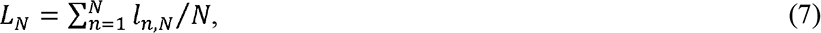

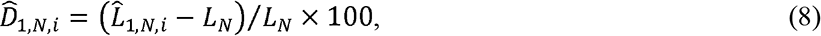

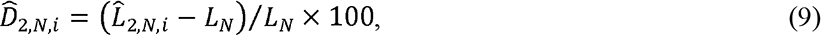

where *L_N_* is the true mean total length in the fish box when the true sample size was *N* and *L_N_* did not change for each shot *i*. A simple regression analysis was conducted where the response and independent variables were *D̂*_1,*N,i*_ and detected rate to confirm if the difference was affected by the number of fish in the box.

## 3. Results

### 3.1 Confusion matrix

Because the loss function from the validation data was minimized at the 32^nd^ epoch, weight parameters were used at that time. The confusion matrix obtained from the dataset used for the validation is presented (Table 3). The precision and recall for “F-100” and “F-other” were 0.91 and 0.61, and 0.87 and 0.63, respectively. Because we only counted classes in which the probability was larger than 0.8, misses were high, especially for “F-other.”

**Table 3.**
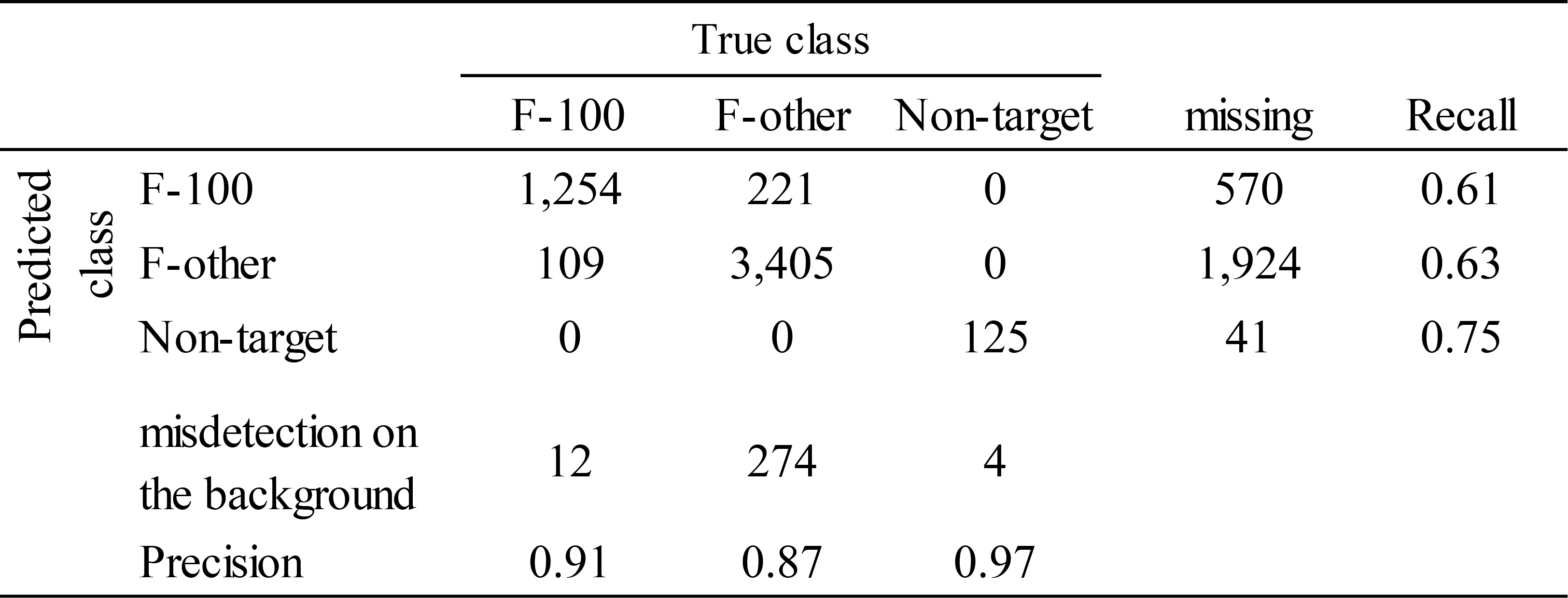
Confusion matrix and calculated precision and recall each class. *Missing* indicates that the fish were there but missed as they were considered as background. *Misdetection on the background* indicates that fish were detected as present, even though there were no fish in the background.

### 3.2 Experiment I

The difference *d̂*_*n*_ between the estimated and measured values obtained for each fish species is shown on the y-axis and the measured value on the x-axis of Fig. 5a–f. 99%, 98%, 92%, 96%, and 97% of *d̂*_*n*_ were included within 5% intervals for mackerels, Japanese jack mackerels, Japanese sardines, red barracudas, and bullet tuna, respectively. In addition, all the bias *B̂* was within ± 3%, where a maximum and minimum bias was 1.15 and -2.77, respectively. The smallest bias was 0.14 for mackerels.

**Fig. 5.**
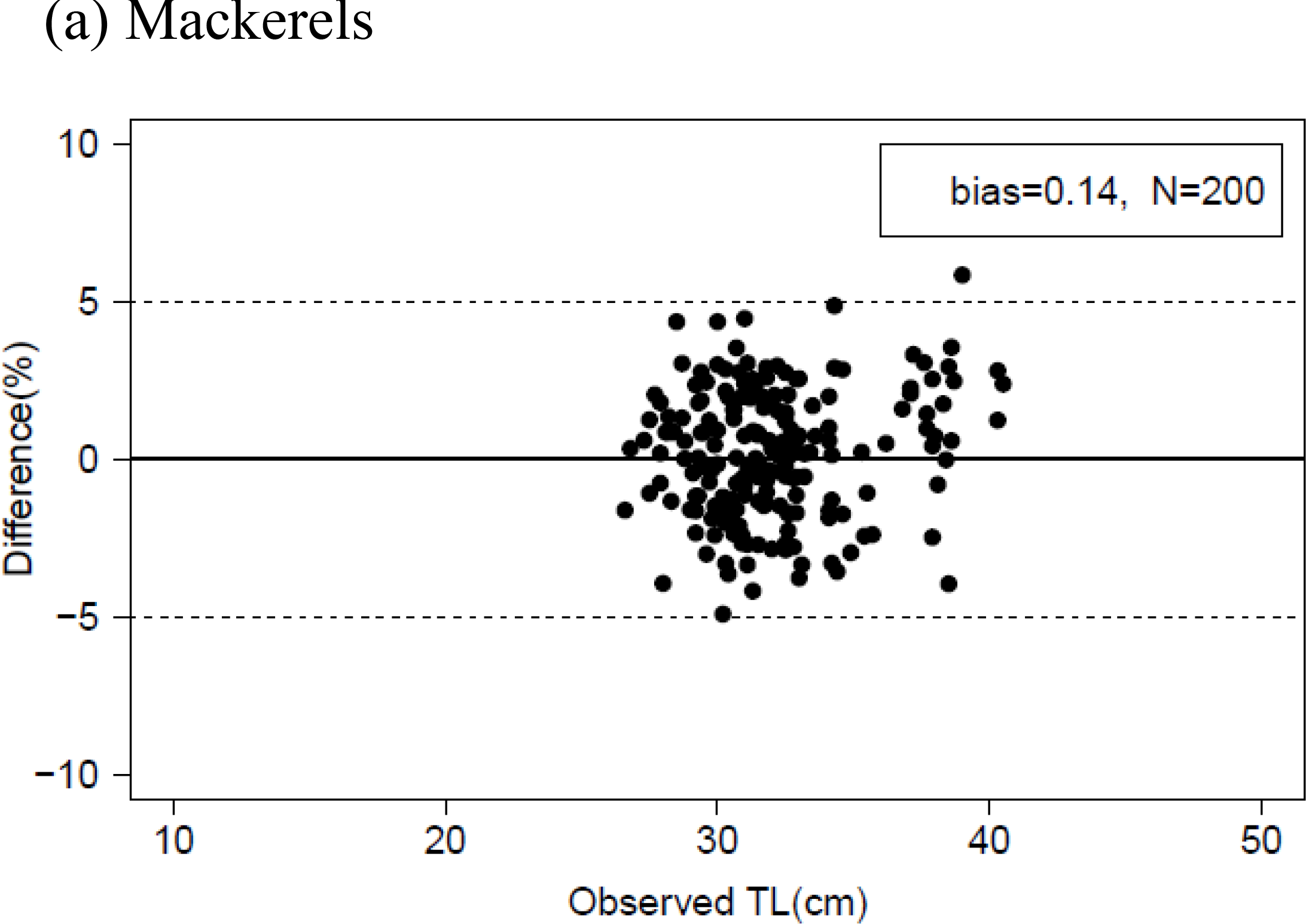
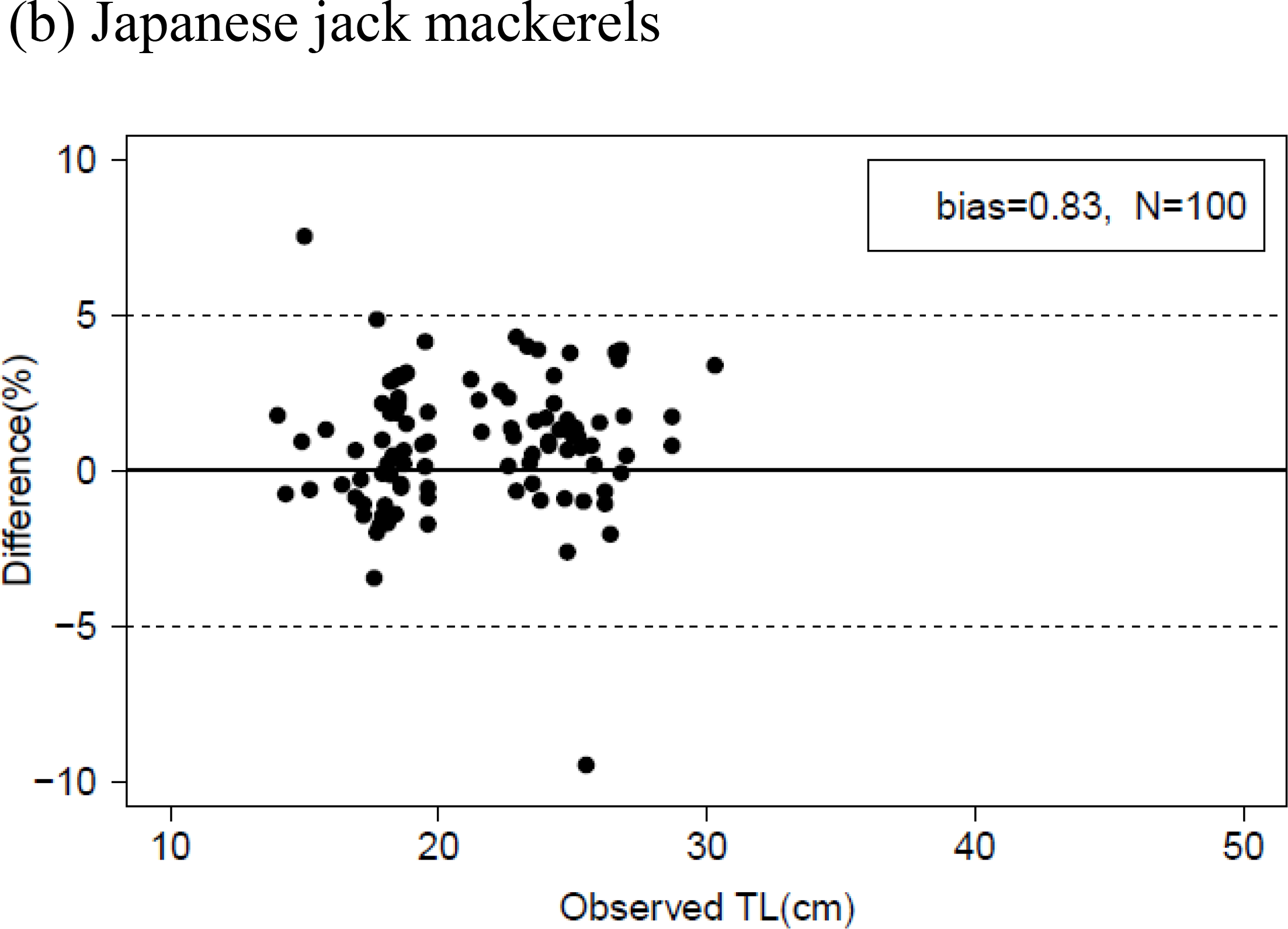
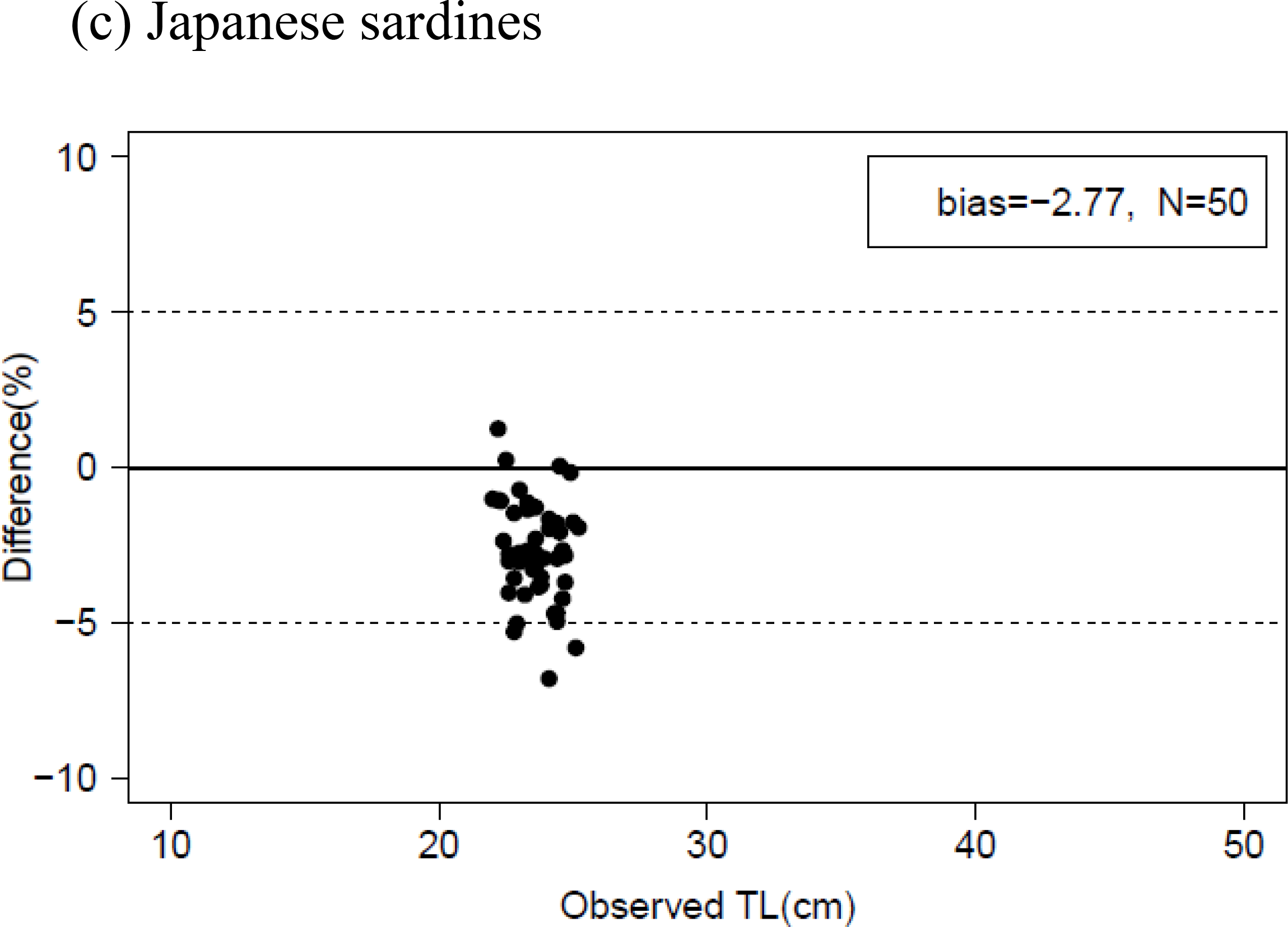
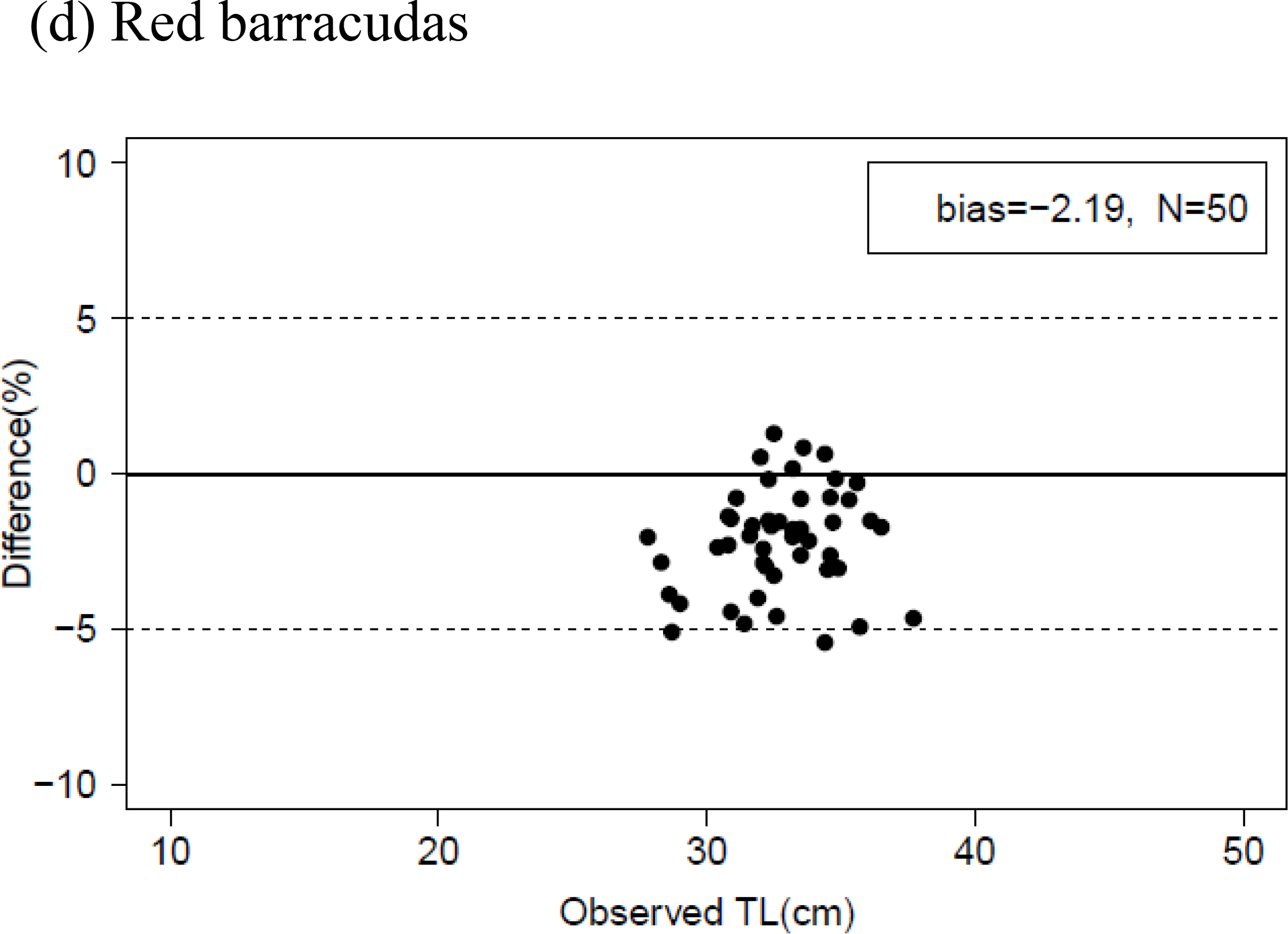
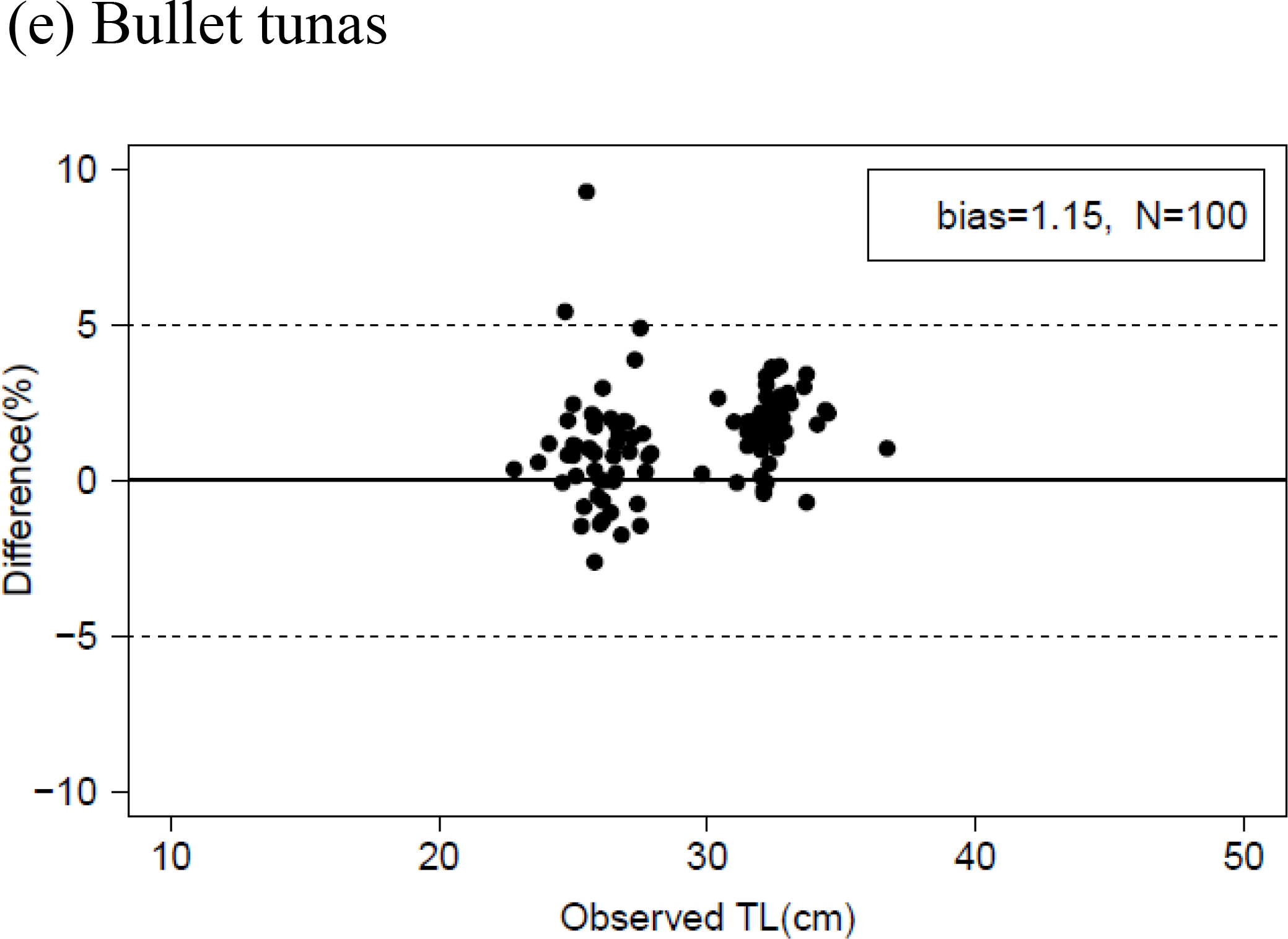
The relative difference *d̂*_*n*_ (%) between the estimates and observed total length (TL) for each fish species: (a) Mackerels, (b) Japanese jack mackerels, (c) Japanese sardines, (d) Red barracudas and (e) Bullet tunas. Black circles indicate the values of the difference. The calculated relative bias *B̂*_*n*_ and sample size are also shown in legends. Black break line shows ±5%.

### 3.3 Experiment II

Detected rates of fish that were identified as both “F-100” and “F-other” decreased as the true number of fish in the box *N* increased (Fig. 6a, b). The rates rapidly decreased for the “F-100” individual, although the rates of “F-other” gradually decreased, and some of them were over 100% because some parts of the fish body were detected twice.

**Fig. 6.**
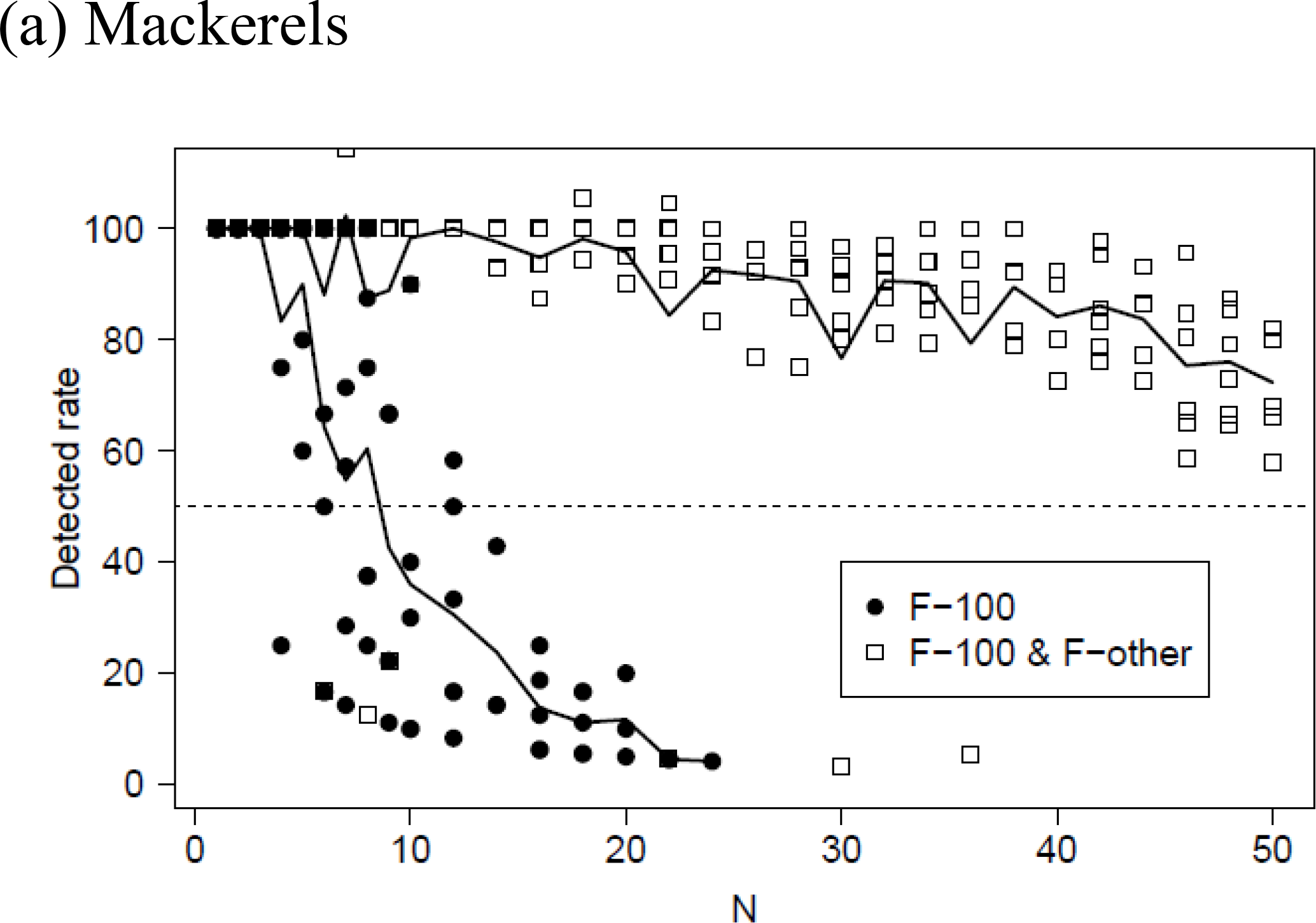
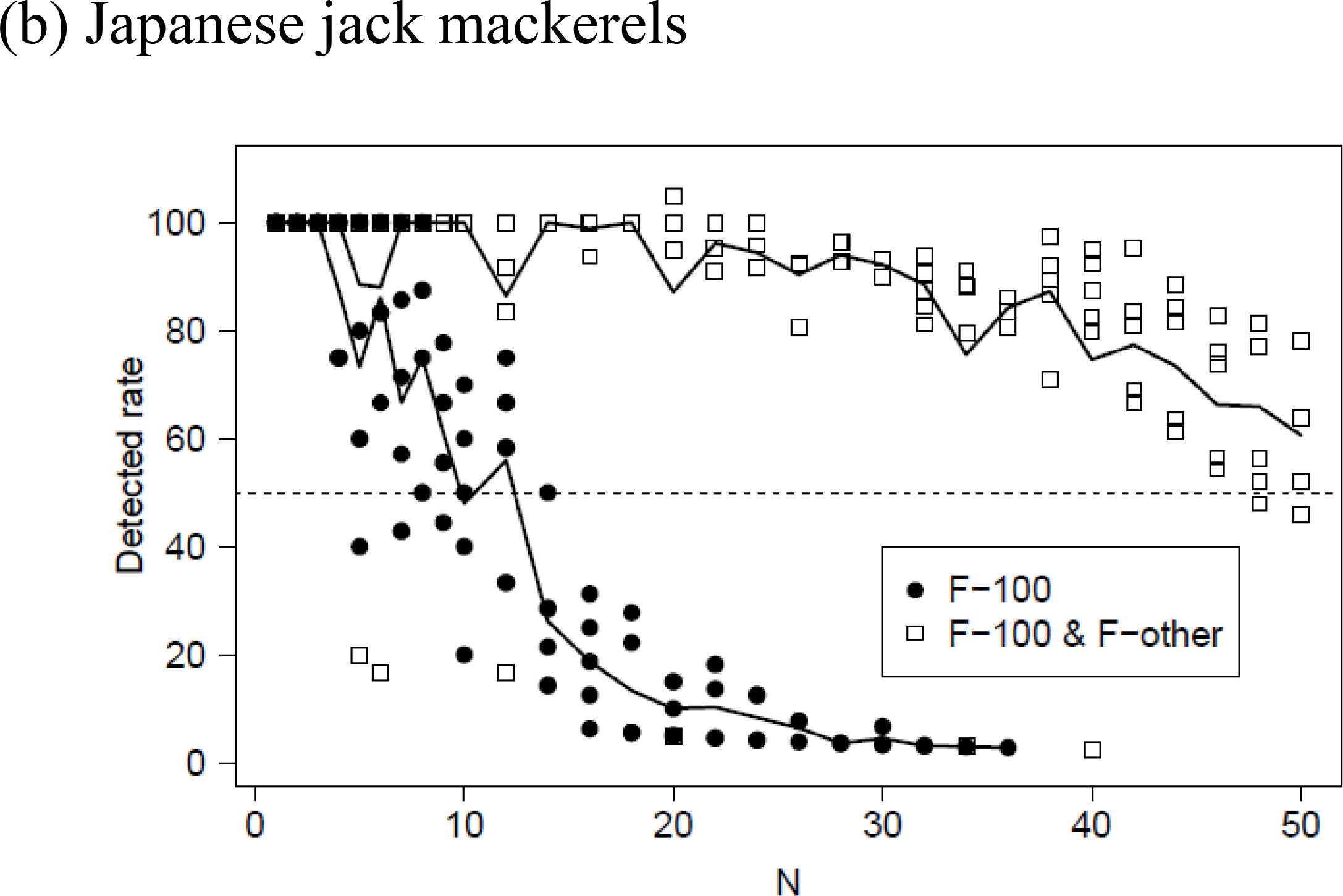
Relationship between the detected rates and the true number of fish in the box (*N*); (a) Mackerels and (b) Japanese jack mackerels. Black circles indicate values of the rate calculated from only non-occluded individuals (*R̂*_1,*N,i*_), and the white squares indicate both non-occluded and occluded (*R̂*_2,*N,i*_) each shot *i*. Black bold and break lines indicate those mean values at each *i* and 50% of the detected rate, respectively.

The relationships between *D̂*_1,*N,i*_, and the detected rates are shown in Fig. 7. The value of *D̂*_2,*N,i*_ took on larger negative values as the detected rate decreased, although it did not decrease when only “F-100” was extracted. The variances of *D̂*_2,*N,i*_ increased when the detected rates were low. The parameters from the simple regression analysis are shown in Fig. 7. The estimates showed that *D̂*_1,*N,i*_ was not affected by the detected rates, and mean differences (i.e., regression line) were included within, at most, *D̂*_1,*N,i*_ 1.5%, where the detected rate (i.e., the independent variable *x*) was changed from 0 to 100. Examples of predicted labels and mask areas for every shot of Japanese jack mackerel, where *N* = 14, are shown in Fig. 8.

**Fig. 7.**
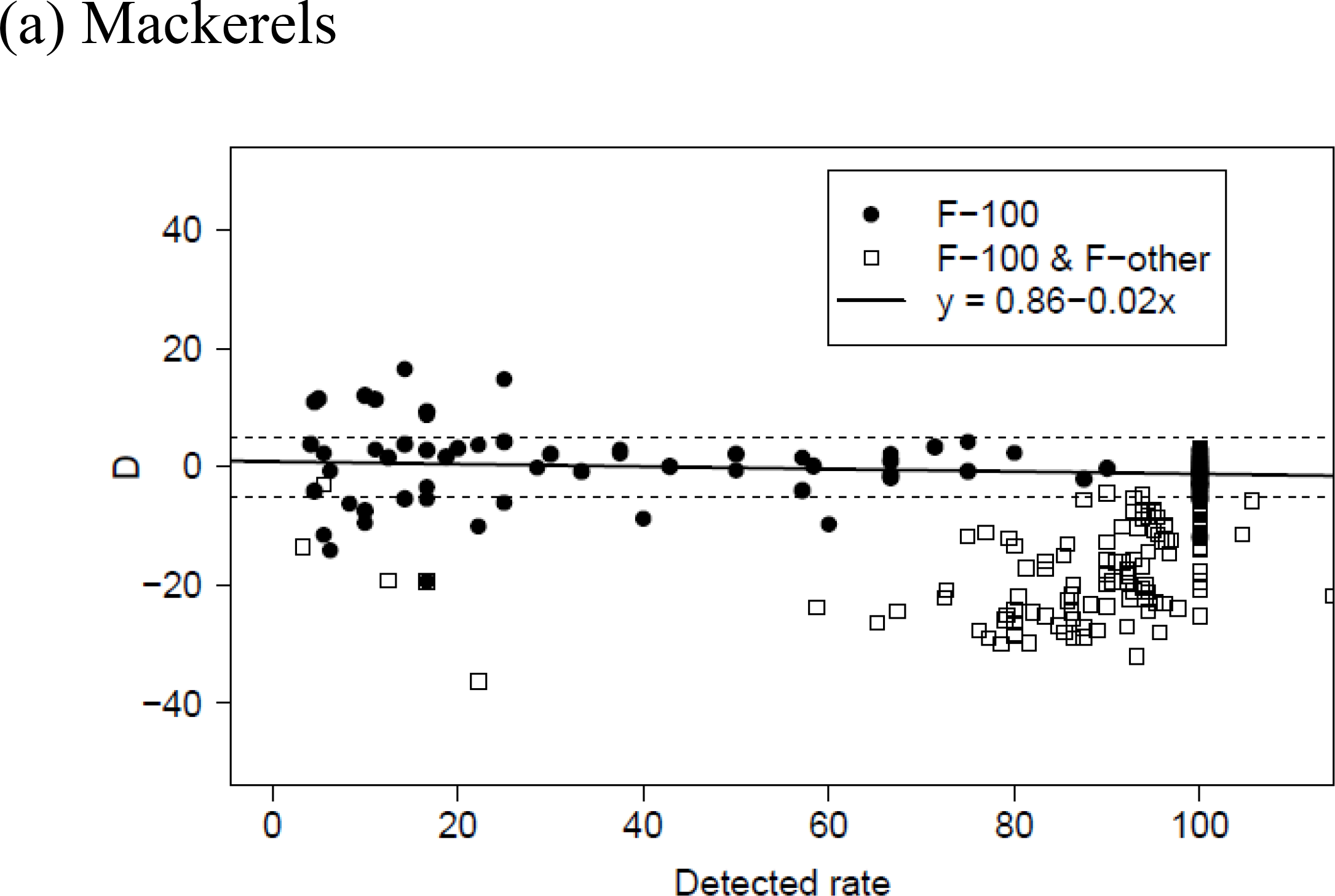
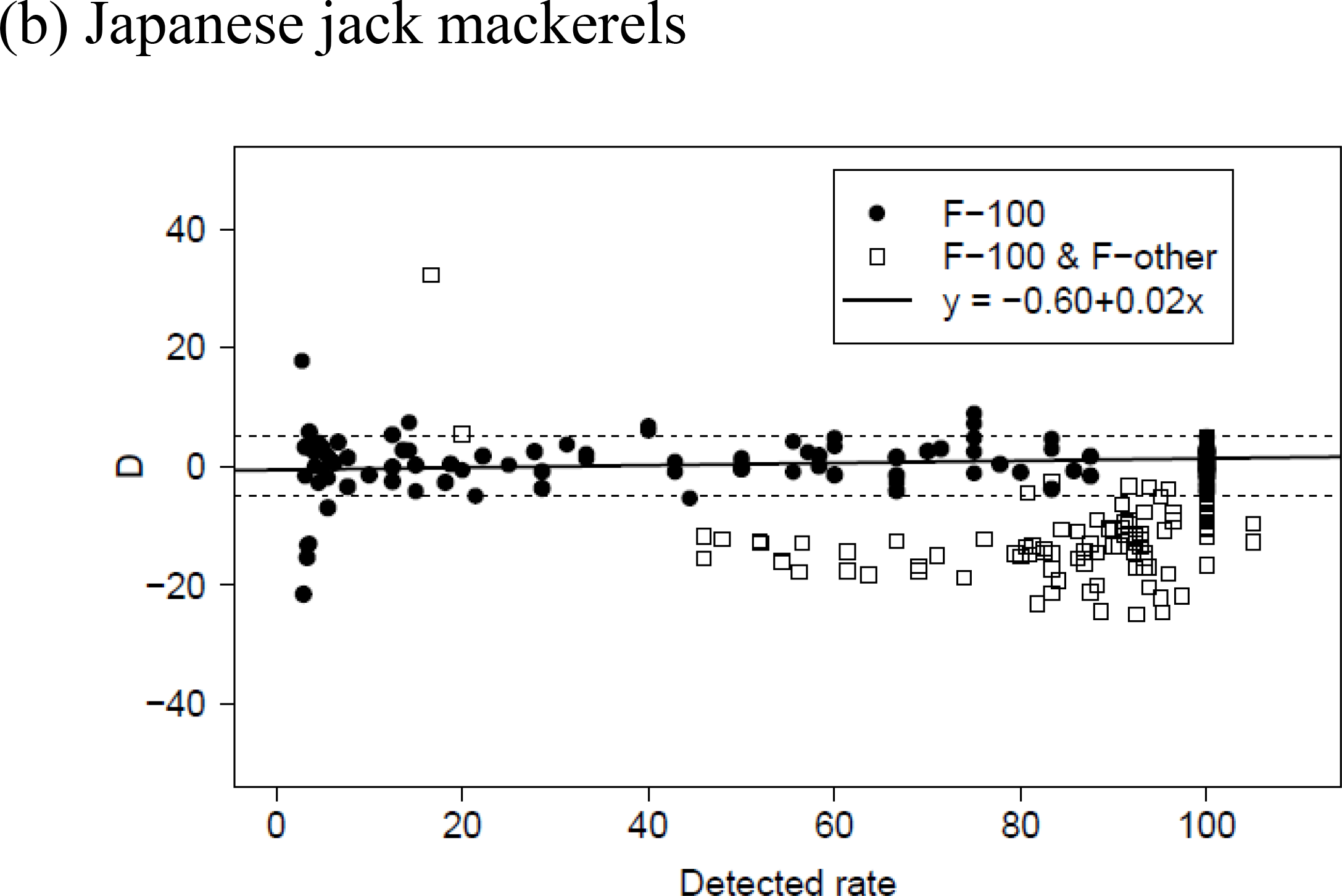
Relationship between the detected rates and mean difference of estimated and observed total length composition; (a) Mackerels and (b) Japanese jack mackerels. Black circles indicate values of the difference calculated from only non-occluded individuals (*D̂*_1,*N,i*_) and the white squares indicate both non-occluded and occluded (*D̂*_2,*N,i*_) each shot *i*. Black break line shows ±5%.

**Fig. 8.**
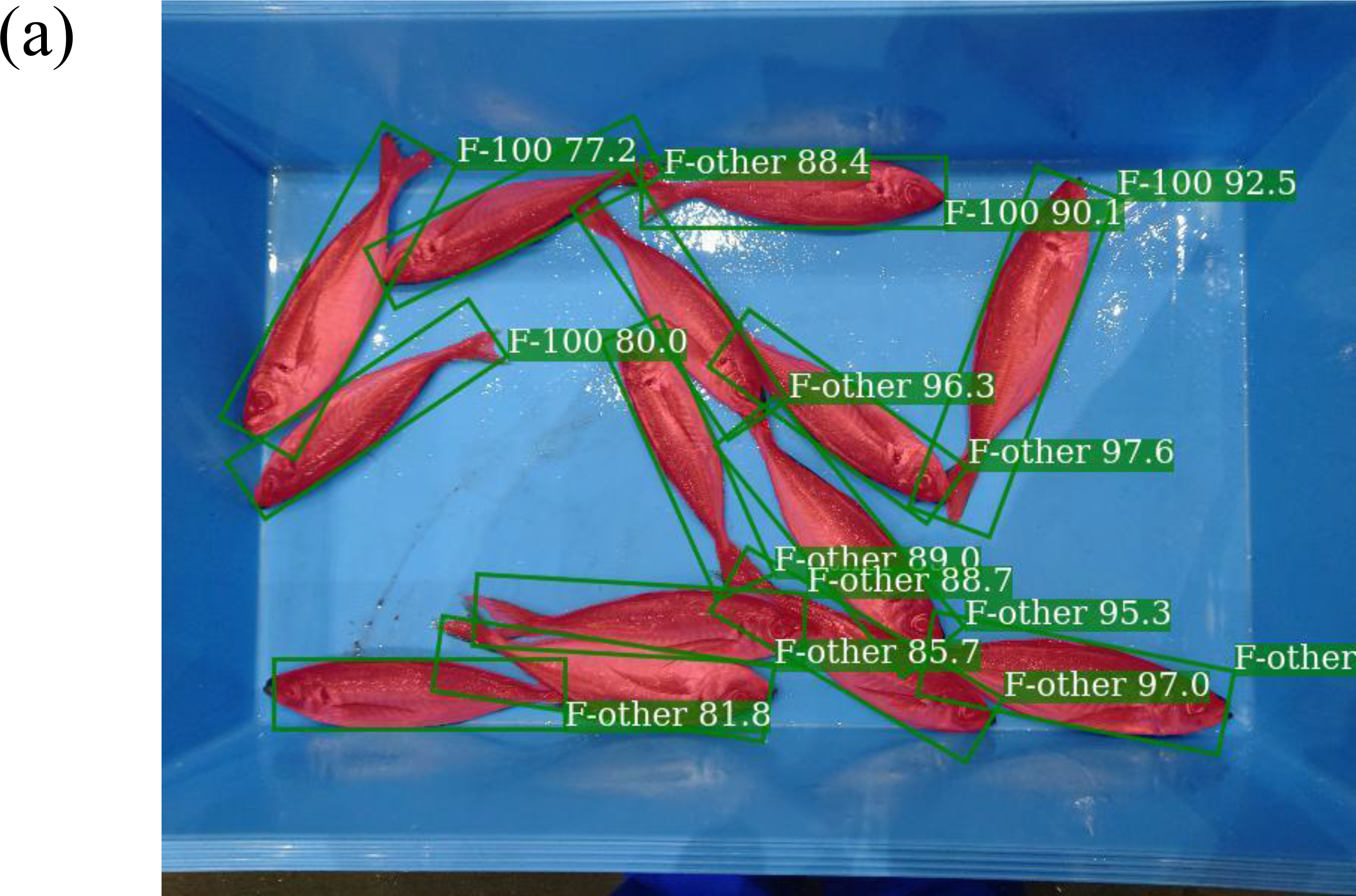
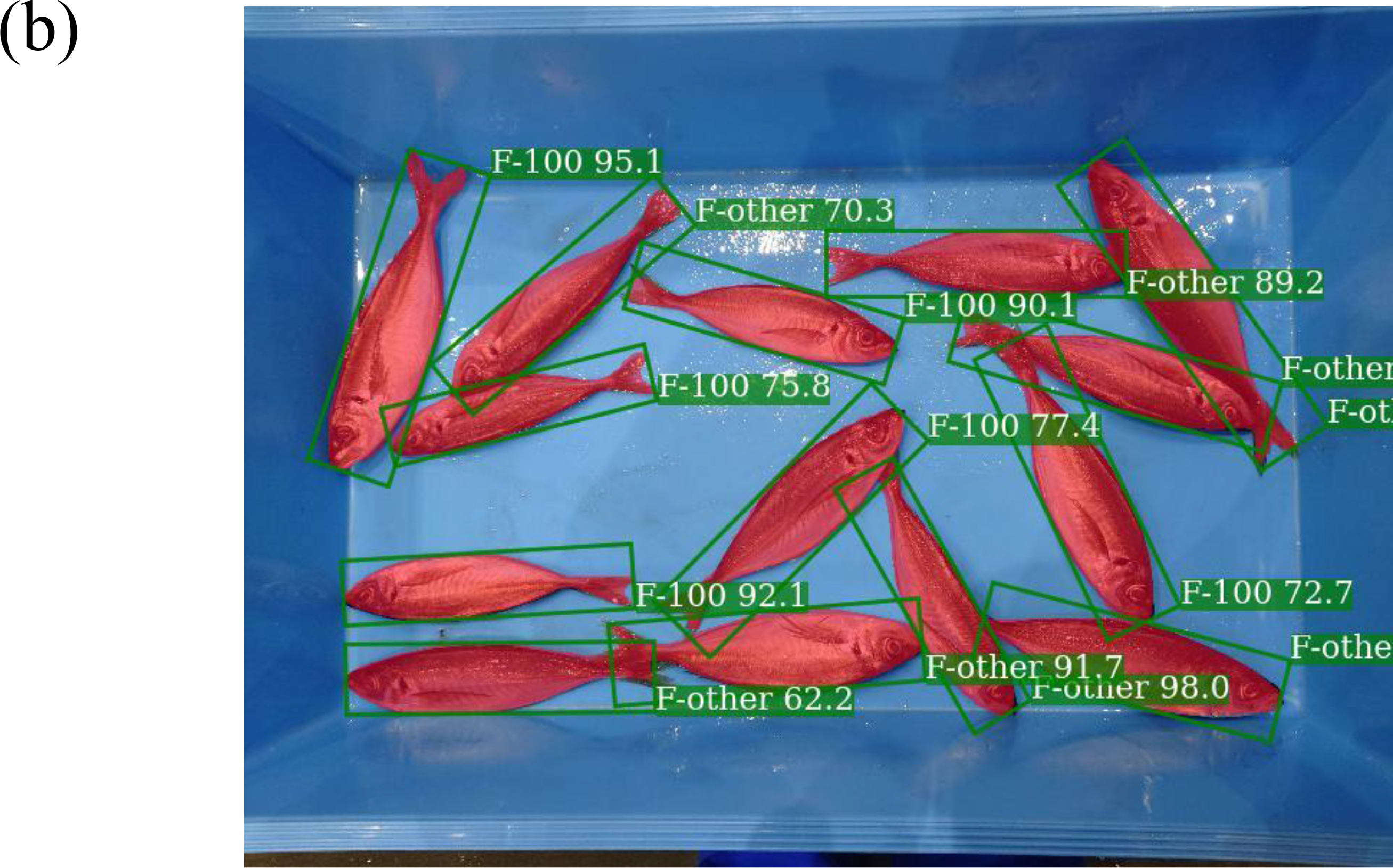
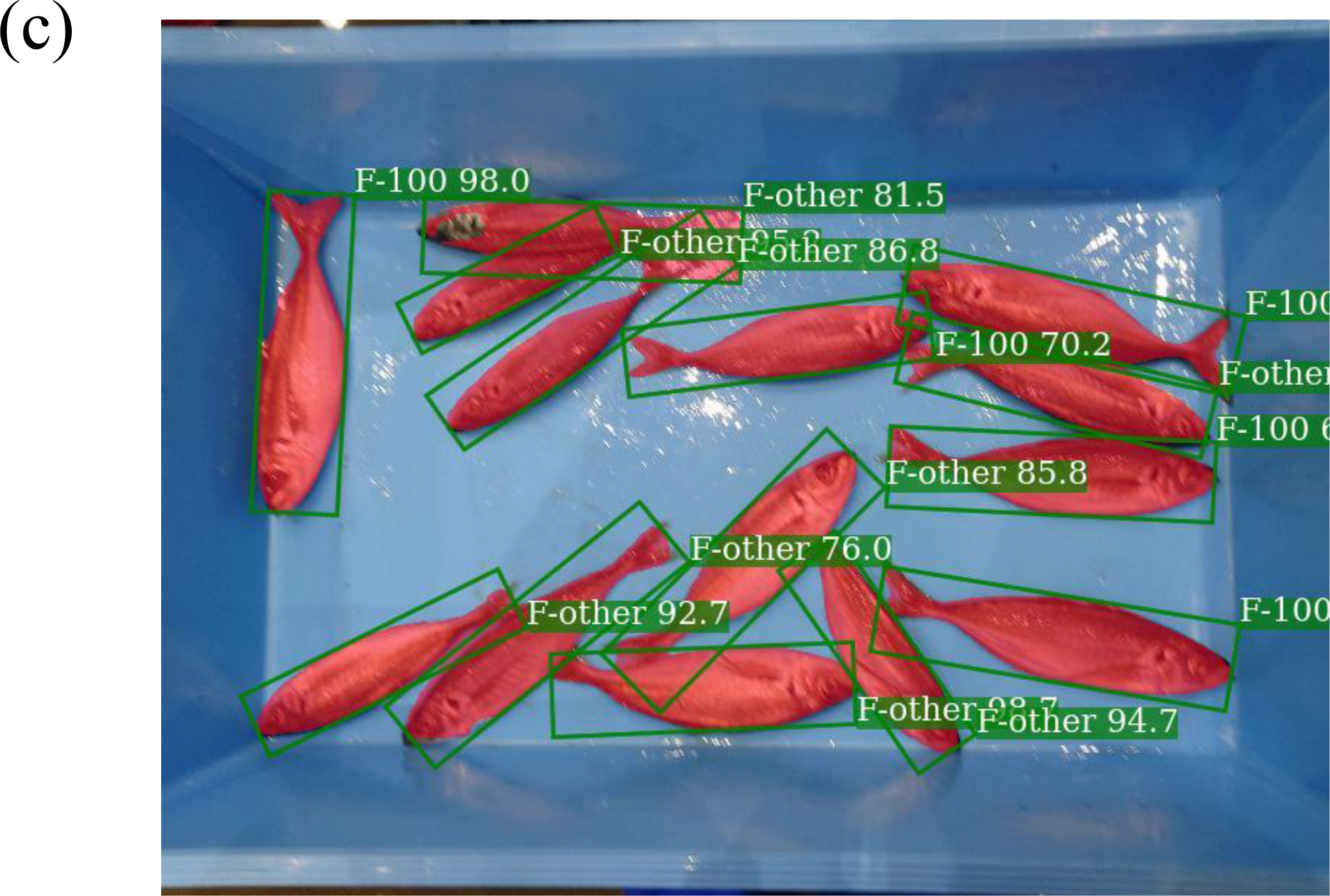
Example of predictions for Japanese jack mackerels (*N*=14). The positions of the fish body is different because it was photographed three times (a: first time, b: second time, c: third time) after being stirred by hand. In the third shot, the image is slightly blurred due to the camera shaking.

## 4. Discussion

Few studies have used deep learning to estimate the body length of fish using 2D images. Previous studies have estimated the total length from the size of a fish’s head (Álvarez-Ellacuría et al., 2020), estimated average body size from the weight of fish in a fish box (Palmer et al., 2022), estimated size directly by deep learning (Ovalle et al., 2022). In all these studies, the camera was fixed and was capable of capturing images of the fish directly below. The distance between the camera and the target fish also does not change from one image to the next; therefore, the value of cm/pixel remains constant from image to image. In our study, we showed that total length estimation is possible even in situations in which the camera cannot be fixed (e.g., no place or high cost to setup, no power supply, and poor security). This will increase opportunities to apply fish length estimation through deep learning, regardless of whether the camera can be fixed.

The detected rate of fish detected as “F-100” decreased rapidly and eventually reached zero (Fig. 6a, b). This is because as the number of fish in the fish box increases, the probability of occlusion increases. This was also supported by the results shown in Fig. 7. Here, the difference between the average length of individuals detected as “F-100” and their actual mean length (i.e., the regression lines) was within ±1.5%, regardless of the detected rate. If the individuals detected as “F-100” included individuals that were underlain by other fish (i.e., “F-other”), the difference would have increased to a negative value as the detected rate decreased. In fact, the mean difference was negative only in the results that included both individuals detected as “F-100” and “F-other.” This was because the total length was calculated from individuals smaller than the actual size as “F-other.” These results suggest that fish detected as “F-100” do not include occluded fish, regardless of the degree of occlusion. In other words, when the total length was estimated only from individual fish detected as non-occluded, regardless of how many fish were placed in the fish box and photographed, the difference from the actual mean length would be within ±1.5% on average, or all would be processed as occluded individuals, which would result in an incorrect estimation of the total length composition. Because it is difficult to completely control the number of fish in the fish box, this fact simplifies the procedure by which a photographer can make length estimates using the proposed method.

There is a trade-off between recall and precision. To obtain an accurate total length composition of the target fish, the recall can be low, but the precision must be high. In this study, when the probability was set to 0.8, 8% (1–0.92) of the fish were misclassified as “F-100”. To obtain the total length composition, it is desirable to maintain a high probability (e.g., score = 0.8). For example, for species that are abundant and frequently photographed, a more stringent probability value would provide accurate total length composition from many images. For species with many stock management constraints, an accurate total length composition is important; therefore, a higher probability is desirable. However, when 99% of the fish body is visible, it is necessary to label the *exposure* as “F-other,” but this is often difficult for annotators to determine. It is very important to determine the judgment criterion of the label in advance to detect “F-100” accurately, even if the probability is high.

If total length estimation can be performed using ToroCam and deep learning techniques, there are two advantages to conducting total length measurements. The first is a reduction in the work time. The total length of catch at fishing ports or the fish market is collected manually by measurers. In most cases, the data were recorded by punching holes in the measurement paper using an eyeleteer. After returning to the laboratory, the positions of the holes in the measurement paper are read and converted to numerical values and entered in a spreadsheet such as Excel. However, the proposed method can significantly reduce the labor hours of the measurers because the total length estimate is output simply by extracting data from a smartphone, and a deep learning model makes the inference. The authors experimentally calculated that the time required to automatically output numerical values from a fish image to a spreadsheet using this method was only 9% of the time required to read the values from the holes in the measurement sheet and type them into a spreadsheet.

The second advantage is that the time required for measurement is reduced, allowing the collection of the total length information for a greater number of fish. In this study, we showed that we could estimate six fish species with an accuracy of ± 3%, and we expect that fish species not considered in this study can also be detected as “F-100” in the same manner if the sample size of the training data is increased. This will update the conventional stock assessment methods by incorporating length-based models (Hordyk et al., 2014), which are expected to improve the accuracy of stock assessments for several fish species.

When a measurer takes a fish image manually with ToroCam, slight movements of the hand would change the position of the rectangle, making it time-consuming for sensitive users to align the four corners of the fish box precisely. However, in practice, the size of the fish box was sufficiently large relative to the image size, and slight shaking did not have a significant effect on the bias. The act of aligning the rectangle with the four corners of the fish box also played a role in enhancing the effect of shooting the fish directly above. It would be good to confirm that the difference between the estimates and true length does not vary greatly from person to person before actually taking the images. As a future issue, linking the transmission function of the smartphone with ToroCam would further facilitate the process of obtaining length compositions.

Future studies should include a combination of fish species identifications. There have been reports of the impact of individual occlusions on fish species classification (Ovalle et al., 2022). In a previous study, the degree of occlusion was manually separated; however, it was suggested that learning the occlusion itself would reduce this effort. Although our study did not classify fish species, it is possible to combine fish species classification models, which will be the next step. In such cases, the burden on measurers should be further reduced.

## 5. Conclusion

In this study, a smartphone application, ToroCam, and deep learning method were used to detect fish that were not occluded. The performance was within ± 1.5%, and reliable total length estimates were obtained. Increasing the amount of stock used within the maximum sustainable yield (MSY) is an international goal described in the Sustainable Development Goals (SDGs). The results of this study will contribute to the calculation of MSY for several fish species that conventional EMS cannot target, because the total length composition is available simply by photographing.

## Acknowledgement

The authors would like to thank Mr. Ken Nakagawa, Mr. Satoshi Kuwahara, Dr. Toru Kitamura, and Mr. Yasuaki Matsumoto for their assistance in obtaining fish images. We would also like to thank Mr. Kazuharu Iwasaki for providing the original code for carrying out the Mask R-CNN. Dr. Yutaka Osada worked with us to acquire images of the fish and provided helpful comments early in the analysis. Moreover, the authors would like to thank Editage (www.editage.com) for English language editing and TTPM Inc. for providing high-quality annotated data. This study was funded by the Fisheries Agency of the Ministry of Agriculture, Forestry and Fisheries of Japan.

---

Supplementary Table 1 Although identification of species names using deep learning techniques was not carried out in this study, the species name had been annotated; otherwise, their genus, family, or order names were annotated if the species name of the fish could not be identified (e.g., their body position was not appropriate to identify). Species for which species identification was not possible for some reason and that were similar in appearance were treated as the same group of fish species, even if their genus, family, and order were different.

